# Targeting LC3/GABARAP for degrader development and autophagy modulation

**DOI:** 10.1101/2023.10.05.560930

**Authors:** Martin P. Schwalm, Johannes Dopfer, Adarsh Kumar, Francesco A. Greco, Nicolas Bauer, Frank Löhr, Jan Heering, Sara Cano, Severin Lechner, Thomas Hanke, Ivana Bekic, Viktoria Morasch, Daren Fearon, Peter G. Marples, Charles W. E. Tomlinson, Lorene Brunello, Krishna Saxena, Nathan Adams, Frank von-Delft, Susanne Müller, Alexandra Stolz, Ewgenij Proschak, Bernhard Kuster, Stefan Knapp, Vladimir V. Rogov

## Abstract

Recent successes in developing small-molecule degraders that act through the ubiquitin system have spurred efforts to extend this technology to other mechanisms, including the autophagosomal-lysosomal pathway. Therefore, reports of autophagosome tethering compounds (ATTECs) have received considerable attention from the drug development community. ATTECs are based on the target recruitment to LC3/GABARAP, a family of membrane-bound proteins that tether autophagy receptors to the autophagosome. In order to validate the existing ligands, we rigorously tested target engagement of reported ATTEC ligands and handles. Surprisingly, using various biophysical methods, most available ligands did not interact with their designated target LC3. Intrigued by the idea of developing ATTECs, we evaluated the druggability of LC3/GABARAP by *in silico* docking and large scale crystallographic fragment screening. The data revealed that most fragments bound to the HP2, but not the HP1 pocket of the LC3-interacting region (LIR) docking site, suggesting favorable druggability of this binding pocket. Here, we present diverse comprehensively validated ligands for future ATTEC development.

## Introduction

Targeted protein degradation (TPD) has received great deal of attention based on the potential of chemical degraders to become a new modality in drug development.^1^ Two major strategies are currently used: molecular glues (glues) and PROTACs (PROteolysis TArgeting Chimeras).^2^ Glues bind to an E3 ligase and recruit a protein of interest (POI) with their solvent exposed moieties. This chemically induced proximity of the POI and the E3 ligase results in ubiquitination of the POI and subsequently in its proteasomal degradation. PROTACs trigger selective degradation by a similar mechanism but they are chimeric molecules using two distinct ligands, one binding to an E3 ligase and one to the POI, connected by an appropriate linker moiety.^3^ PROTACs and glues have hugely expand the druggable target space as they can bind anywhere to a POI and not only to a specific binding site relevant to disease development. Additionally, their properties of acting catalytically and often degrading the POI highly selectively bears a promise that these new drug modalities could be effective at very low compound concentrations reducing drug toxicity.^4^ Spawned by the potential of selective degraders in drug development, new pathways have been explored to extend the toolbox that can be utilized for the design of these molecules. Among them are LYTACs^5^ (LYsosome-TArgeting Chimeras) for the degradation of membrane proteins as well as ATTECs^6^ (AuTophagosome TEthering Compound) which hijack the autophagy/lysosomal pathway for selective degradation of POIs. Excitingly, these ubiquitin independent systems would also allow degradation of large organelles, pathogens and protein complexes.

Macro-autophagy (Autophagy hereafter) is a fundamental cellular process regulating degradation and recycling of cellular components,^7, 8^ also allowing the removal of bulky cytosolic cargo, such as large protein complexes, lipid droplets, portions of and whole organelles, and even bacteria that invaded the cytoplasm.^9, 10^ Cargo degradation is achieved by enclosure into a double-membrane vesicle (autophagosome) followed by autophagosome trafficking to the lysosome for degradation. Autophagy is an evolutionarily conserved complex process orchestrated by ∼40 autophagy-related (Atg) proteins, which include, among others, the autophagy-related ubiquitin-like modifiers (Atg8 in yeast) LC3A, LC3B, LC3C, GABARAP, GABARAPL1, and GABARAPL2 proteins (LC3/GABARAP hereafter).^11^ LC3/GABARAP recruit cargo-receptor-complexes by a short sequence motif called the LIR (LC3-Interacting Region), mediating autophagosomal recruitment and degradation by interaction with the LDS (LIR docking site) (Figure 1, left plot).^12-14^ In addition to the LDS, LC3/GABARAPs possess an additional interaction site located at the opposite face of the LDS (Figure 1A, right plot), reminiscent of a hydrophobic binding patch present in ubiquitin. Accordingly, this site binds to several ubiquitin-interacting motifs (UIM) and was therefore named UIM docking site (UDS).^15^ Similar to the E3 ligase dependent TPD, small molecules binding to LDS and the UDS probably interferes with cargo recruitment to LC3/GABARAP proteins and could be developed into small molecule degraders by recruitment of targets to the autophagosome.^16^ However, discovery of potent LC3/GABARAP ligands has remained challenging, possibly due to the conformational plasticity of LC3/GABARAP resulting in at least partial occlusion of the LDS.^17^ First, LC3A/B targeting reversible ligands such as the antibiotic Novobiocin^18^, covalent lysine targeting ligands^19^ as well as a number of low molecular weight fragments (overviewed in Figure 1C) have been described binding to the LDS,^20^ but no highly potent ligands or ligands for the four remaining LC3/GABARAPs have been described. Interestingly, in 2019, first ATTECs have been reported suggesting that the autophagy degradation pathway can be exploited for the design of selective degrader small molecules.^6^ In this study, the authors presented a mechanism, where small molecules mediated the autophagosomal degradation of mutant huntingtin protein (mHTT) by LC3A/LC3B recruitment. Subsequently, the identified compounds (10O5/compound **4** and 8F20/compound **3**) have been used as LC3 ligands for ATTEC design, targeting diverse proteins including the bromodomain containing 4 (BRD4)^21^ and nicotinamide phosphoribosyltransferase (NAMPT),^22^ suggesting a broad utility of these ligands for degrader design. However, as ATTEC development is still in its infancy, no tool compounds such as autophagy pathway inhibitors are available that could serve as controls for the proposed degradation mechanism. In addition, we realized that thorough biophysical characterization, evaluation, cellular target engagement or cell-based controls for the developed ATTEC ligands was largely lacking. Given our interest in the development of new degrader molecules and the autophagy pathways, we rigorously evaluated current LC3 ligands (SI Figure S1) by biophysical binding assays *in vitro* as well as in cell lysates. Surprisingly, we failed to detect any interaction for some of the published LC3 ligands using an array of assay systems suggesting that these ligands potentially act through alternative mechanisms. Intrigued by the concept of hijacking the autophagosomal pathway through target recruitment to LC3/GABARAP, we broadly evaluated the druggability of the LDS and UDS by *in silico* screening of an in-house compound library followed by biophysical validation as well as by high throughput crystallographic fragment screening. The campaigns revealed good druggability of the HP2 site within the LDS, a shallow binding pocket interacting with hydrophobic residues in the LIR motif. Additionally, poor accessibility of the HP1 site interacting with aromatic residues in LIR motifs and initial ligands targeting for UDS binding site were found, which natural binding partner still remain understudied. Our data not only prove druggability of all LC3/GABARAPs, but also present a strategy for the development and evaluation of LDS and UDS ligands as starting points for future ATTEC development.

**Figure 1.**
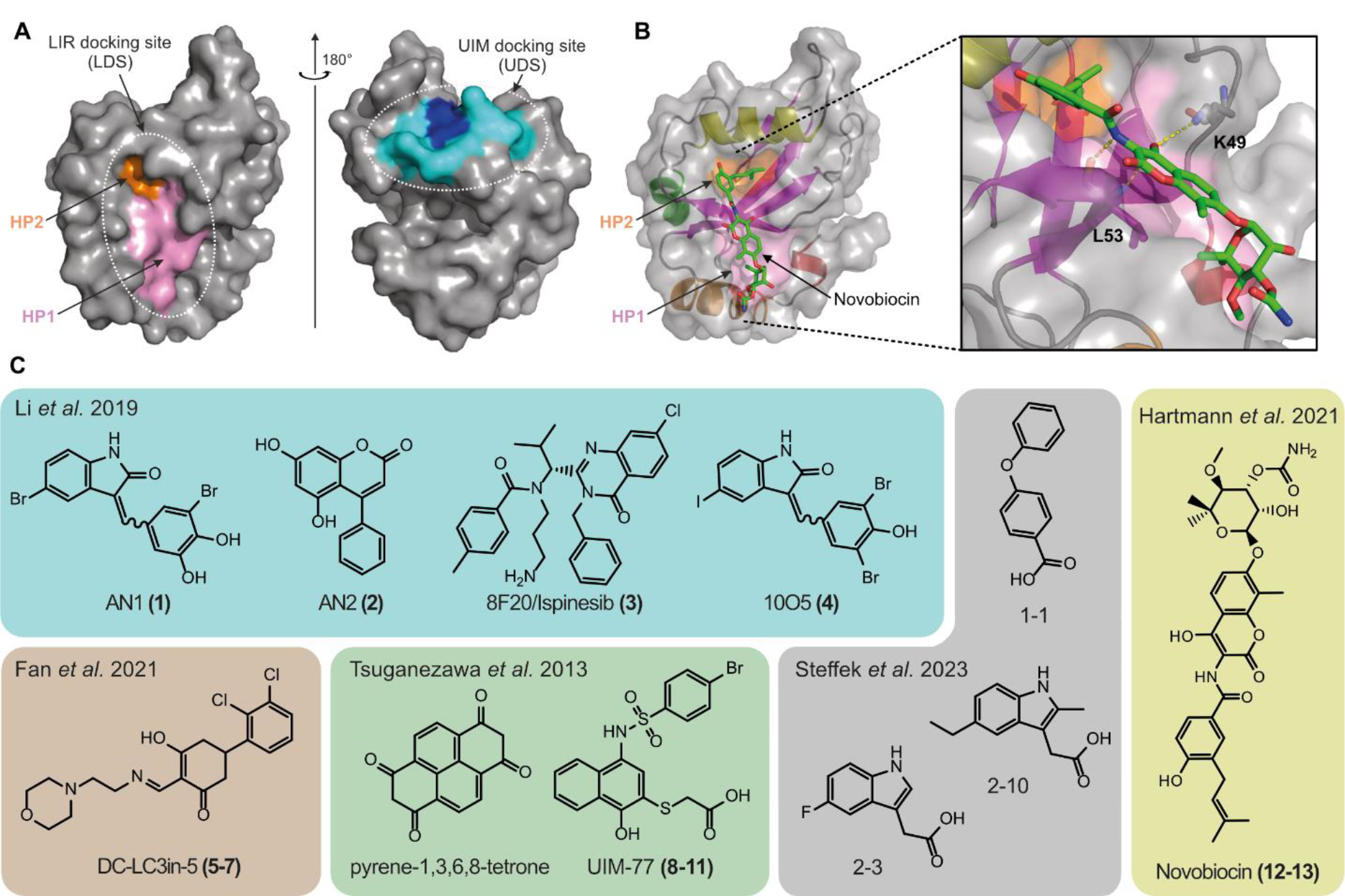
Structural organization of human Atg8 family proteins and reported binder. **A)** left panel depicting the LIR docking site (LDS) of LC3A comprised of hydrophobic pocket 1 (HP1) in pink and hydrophobic pocket 2 (HP2) in orange. Right panel displaying the UIM docking site (UDS) in blue/cyan (pdb: 3ECI). **B)** Dihydronovobiocin bound to LC3A with an enlarged panel of the binding interface by interactions towards K49 and L53 and binding to the HP2 via hydrophobic interactions (pdb: 6TBE). **C)** Chemical structures of compounds, published to bind to LC3/GABARAPs with exemplary structures shown with all structures depicted in SI Figure S1.

## Results

To assess target engagement of reported LC3 ligands, we carried out diverse biophysical binding assays, including fluorescence polarization (FP), isothermal titration calorimetry (ITC) and nuclear magnetic resonance (NMR). We compiled a comprehensive set of ligands reported in recent literature for this comparative interaction study, including all ligands used for ATTEC design (AN1 (**1**), AN2 (**2**), 8F20 (**3**) and 10O5 (**4**))^6^, covalent ligands that target the amine of K49 within the LDS (compounds **5-7**)^19^ and four analogs of ligands and fragments that have been published to disrupt the p62-LC3 interaction (compounds **8-11**).^23^ Additionally, we included Novobiocin (**12**) and Dihydronovobiocin (**13**), which we reported previously as a ligands of LC3A and LC3B.^18^ We also included five LIR peptides spanning a wide affinity range as positive controls. A full list of selected LC3 ligands has been compiled in supplementary SI figure S1 and representative ligands as well as the targeted ligand pockets are shown in Figure 1.

Initially, we used temperature shift assays as a binding assay to evaluate small molecule interaction with the LC3/GABARAP family. However, recorded temperature shifts were relatively small, including data measured for control peptides and we therefore deemed this assay as not suitable for the detection of LC3/GABARAP ligands (SI Figure S2 A). Next, we established an FP assay utilizing the p62-LIR peptide linked to a Cy5 fluorophore as a tracer molecule. Dose dependent titrations using all LC3/GABARAPs yielded assays with good signal to noise ratio and it resulted in measured K_D_ values for the tracer between 3 and 17 μM across the human Atg8 family. Thus, this tracer allowed establishment of a displacement assay suitable for screening and binding affinity determination of ligands in the low micromolar K_D_ range (SI Figure S2 B). We evaluated the established set of ligands (**1**-**13**) against all LC3/GABARAPs with a representative data set for LC3B shown in Figure 2 A and all data in SI Figures S2 C and D. Consistent with data published previously, compound **12** (Novobiocin) bound to LC3A and LC3B with highest affinity K_I_ values of 17.4 and 48.4 μM, respectively (Figure 2B). Next, we focused on the ligands used widely for ATTEC development (**1-4**). The dose dependent titrations against LC3/GABARAP family members are shown in Figure 2 C. Surprisingly, no detectable binding to the published targets LC3A and LC3B was observed for all four ATTEC handles up to a concentration of 100 μM. However, weak interaction was detected for 10O5 (**4**) binding to GABARAPL2 but not to its designated target LC3B. Due to the bright color of compounds **1** and **4** and the discrepancy to literature data, we validated these results further by direct binding assays using 2D NMR titration experiments with ^15^N labelled LC3B protein (Figures 2D and 2E, left plots). This technique does not rely on the competition of a tracer peptide and enables the detection of allosteric LC3 binders. In agreement with our FP data, binding of Novobiocin (**12**) caused large chemical shifts perturbations (CSP) within the LIR binding pocket (Figures 2D and 2E). However, none of the compounds **1-4** resulted in significant CSP even at higher compound concentrations in agreement with our FP binding data. Analysis of the small CSP HN resonances in the backbone, induced by **4** revealed that they are predominantly within HP2 with high estimated K_D_ values of ≥ 200 μM for LC3B (SI Figure S3).

**Figure 2:**
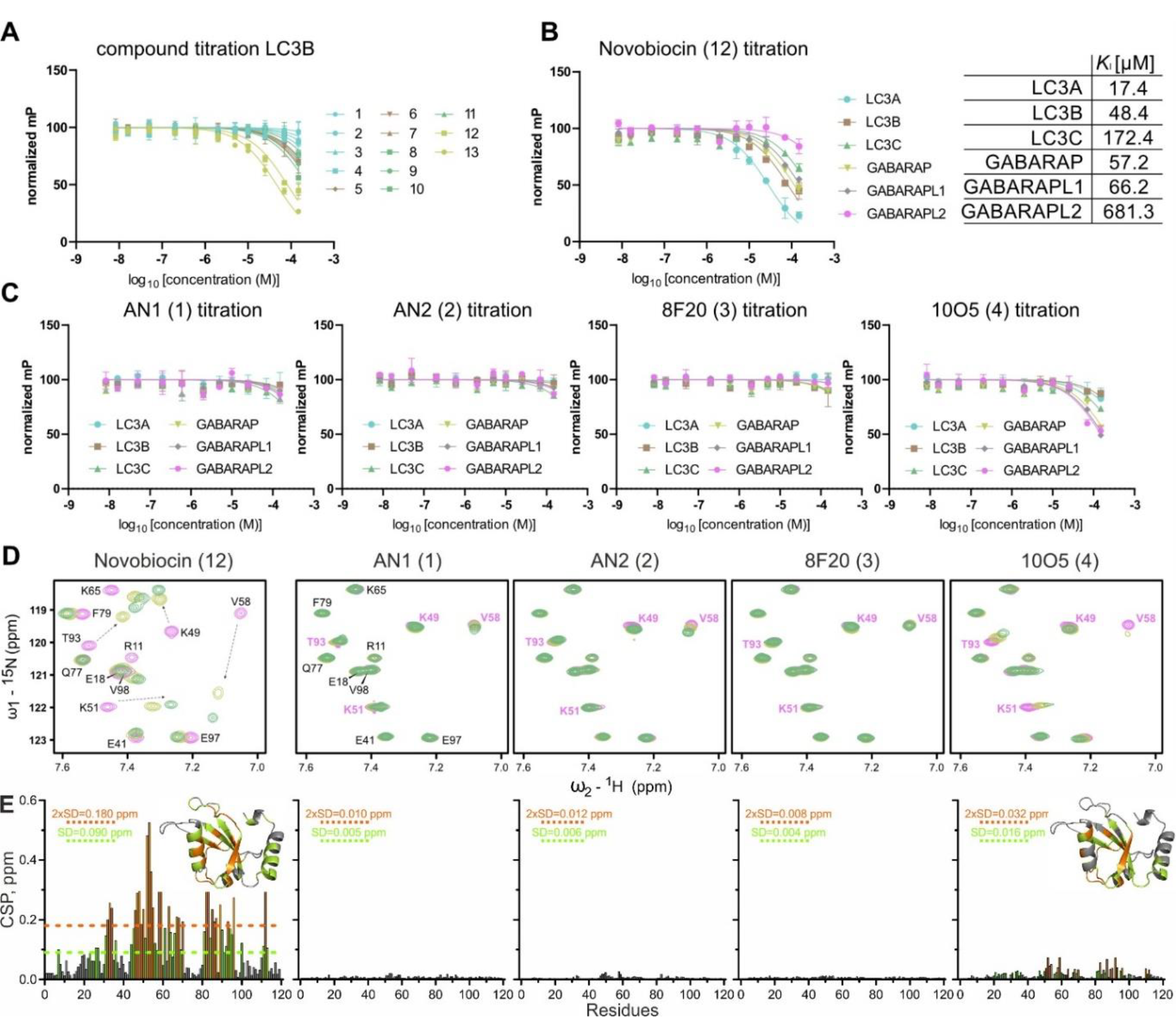
Biophysical characterization of compound-LC3/GABARAP interactions. **A)** and **B)** Fluorescence polarization assay curves obtained from the measured compounds **1**-**13** on LC3B (**A**) and for Novobiocin on all six LC3/GABARAP proteins (**B**) with all curves against all LC3/GABARAPs depicted in SI Figure S2. Assays were run in technical duplicates (n=2) with error bars expressing the SD. Binding curves for all compounds and proteins are depicted in SI Figure S2 **C)** Fluorescence polarization assay curves for compound **1**-**4** against all LC3/GABARAPs. Assays were run in technical duplicates (n=2) with error bars expressing the SD. **D)** Interaction between LC3B and compounds **1**-**4** investigated by NMR. Representative fingerprint areas around the key K51 and V58 backbone HN resonances of [^15^N,^1^H] BEST-TROSY spectra for free LC3B (magenta) and LC3B containing control compound **12** and compounds **1**-**4** at 1:1 (yellow) and 1:2 (green) molar ratios are shown in overlay. Mapping of backbone HN resonances on LC3B sequence and structure are depicted in SI Figure S3. **E)** Left plot: chemical shifts perturbations (CSP) values, induced by **12** at molar ratio 1:2, are plotted against LC3B residue numbers and mapped on 3D-structure (insert). The light green dashed line indicates the standard deviations (SD) over all residues, the orange dashed line indicates double SD values; residues with small (CSP < SD), intermediate (SD < CSP < 2xSD) or strong (2xSD < CSP) CSP values are marked in grey, light green and orange, respectively. Residual plots: compounds **1**-**3** induce insignificant CSP values at molar ratio 1:2, compound **4** induces small CSP around LC3B residues forming HP2 (right plot and insert).

We therefore performed further optimized biophysical analyses, using these methods for LC3A interaction with **1**-**4**. To measure compound interaction via MST, we chose cysteine labelling in order to avoid lysine labeling due to the presence of these residues in the binding sites. After successfully setting up the assay, we were not able to reproduce the published binding data where lysine labeling was used in combination with high protein concentrations of 500 nM (typical range: 5-50 nM) (SI Figure S4 A).^6^ Next, ITC was used as fluorescence-independent method for binding verification. Here, in agreement with earlier experiments, Novobiocin (**12**) revealed binding with a K_D_ (6.7 μM for LC3A), while titrations with AN2 (**2**) and 8F20 (**3**) did not yield significant binding heats. Additionally, we also investigated binding of AN1 (**1**) and 10O5 (**4**) as well as compound (**8**) by ITC but these ligands induced protein precipitation, rendering ITC K_D_ determination impossible (SI Figure S4 B). To reproduce direct binding through FP as reported, we synthesized 8F20 (**17**)- and 10O5 (**16**)-based dye-linked tracer molecules utilizing the same linker attachment point as for ATTEC development (SI Figure S4 C).^22^ Using the reported experimental setup for establishing an FP assay, we successfully reproduced the tracer-LC3 interaction. However, we were unable to obtain displacement data using the parent compound, a standard control in FP assays. The only experimental difference in our FP assay setup was the addition of 0.05 % Tween-20, which is routinely used^24^ to suppress unspecific binding of proteins. Under these conditions, we did not detect any binding, indicating unspecific tracer-LC3A interaction, whereas without Tween-20 some (unspecific) binding was observed (SI Figure S4 D).

Even though we did not observe any binding of compounds **1-4** to LC3 proteins, we were able to measure weak interaction of 10O5 (**4**) with GABARAP family members (Figure 2C). These data motivated us to further investigate interaction to GABARAPL2 by 2D NMR, which confirmed interaction in the HP2 fingerprint area depicted in Figures 3A and 3B (full analysis is shown in SI Figure S5). Indeed, we confirmed weak GABARAPL2-10O5 (**4**) interaction by NMR titration experiments with estimated K_D_ values in the 15-30 μM K_D_ region. Due to the weak interaction with this Atg8 family member, it is not likely that the observed degradation of mHTT was mediated by binding of 10O5 (**4**) to GABARAPL2.^6^ However, due to the use of cell lysate and therefore a lack of cellular and membrane environment contribution factors, we cannot absolutely exclude proteome-wide interactions using this method.

**Figure 3:**
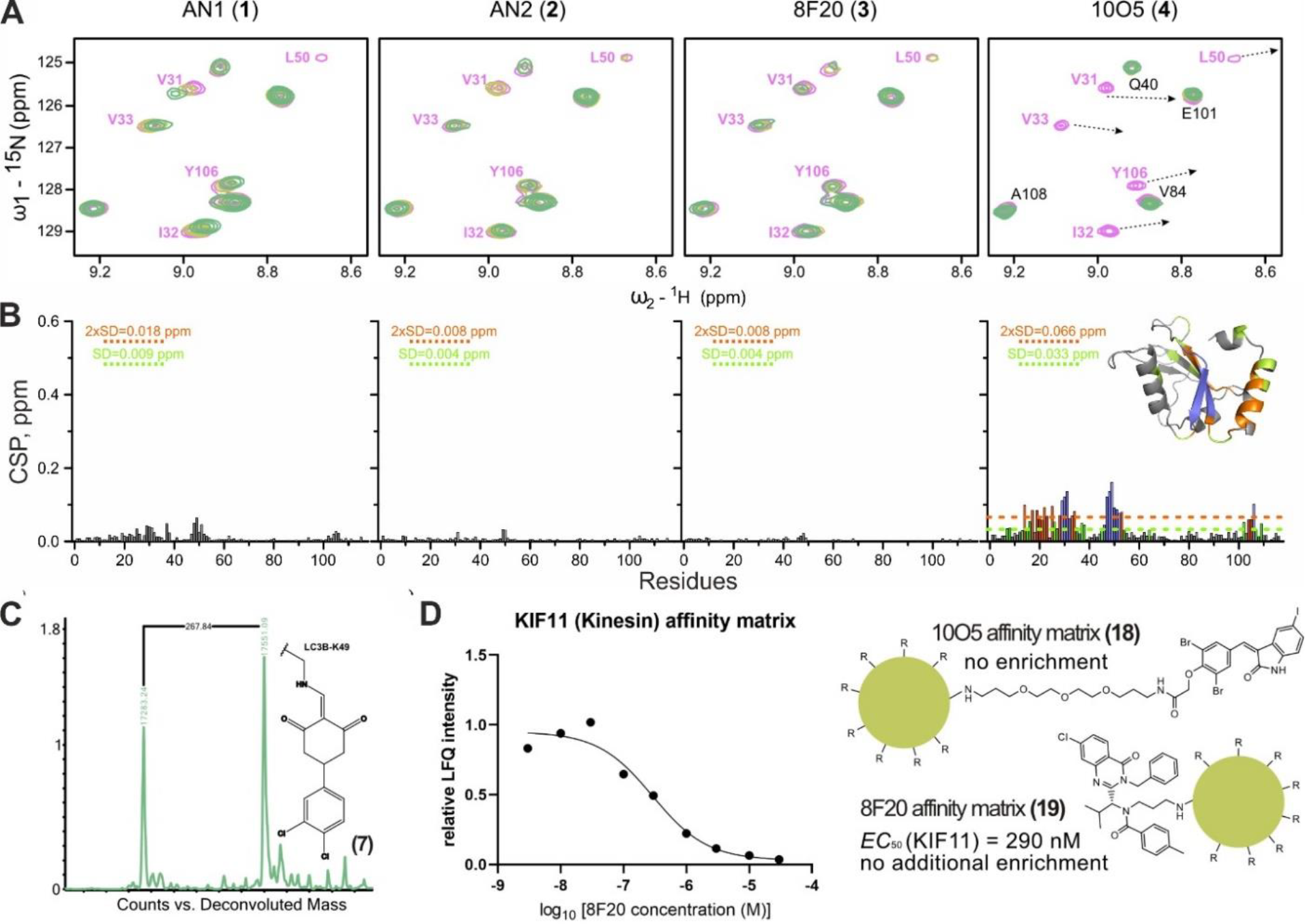
A) Interaction between GABARAPL2 and compounds **1**-**4** investigated by NMR. Representative areas of GABARAPL2 [^15^N,^1^H] BEST-TROSY spectra around the key residues L50, I32 and Y106 backbone HN resonances are shown in overlay with free GABARAPL2 (magenta), and in presence of 1:1 (yellow) and 1:2 (green) molar ratio of each compound (indicated above each plot). Arrows in the plot for GABARAPL2-**4** interaction show directions of large chemical shift perturbations for the resonances which are in the intermediate exchange mode and could not be tracked until the latest titration steps, indicating the strongest interaction of these residues with GABARAPL2. Mapping of backbone HN resonances on GABARAPL2 sequence and structure and additional NMR data analysis are depicted in SI Figure S5. **B)** CSP values, induced by compounds **1-4** at molar ratio 1:2, are plotted against GABARAPL2 residue numbers and mapped on 3D-structure (insert). The light green dashed line indicates the standard deviations (SD) over all residues, the orange dashed line indicates double SD values; residues with small (CSP < SD), intermediate (SD < CSP < 2xSD) or strong (2xSD < CSP) CSP values are marked in grey, light green and orange, respectively. The blue color for sequence- and 3D-mapping for compound **4** are for GABARAPL2 residues which undergo strong intermediate exchange mode (significant decrease of the resonances intensity upon titration with **4**). **C)** Exemplary mass spectrometry data expressing a mass shift of LC3B after treatment with compound **7**. Full data set for compounds **5**-**7** on all LC3/GABARAPs is depicted in SI figure S6. **D)** Chemoproteomic competition assays for target deconvolution of 8F20 (**3**) and 10O5 (**4**). Affinity matrices for target pulldown were generated via amide-coupling, generating (**18**) and (**19**) with NHS-activated sepharose beads. Competition was performed with free **3** and PEG-linked **4** at nine doses and residual binding was calculated relative to a DMSO control. Experiments only identified KIF11 as target (EC_50_ = 290 nM) of **19** while **18-**based competition assay did not lead to enrichment of any targets.

Covalent ligands **5**-**7** only showed weak interaction with LC3/GABARAPs in our FP assay and due to the irreversible nature of this interaction, binding of these ligands might be strongly time dependent. We therefore evaluated the ligands **5**-**7** by ESI mass spectrometry for all LC3/GABARAPs. In agreement with published data,^19^ we detected a mass shift corresponding to the compounds bound to LC3B (Figure 3C). The mass shift corresponded to a single modification, supporting the literature data, suggesting that compounds of this class selectively form a covalent bond to LC3 proteins to K49.^19^ However, treatment of either of the six proteins with compounds **5**-**7** revealed no selectivity within the human Atg8 family members, raising the possibility of further off-targets within the proteome, based on the reactivity of these compounds (SI Figure S6). To investigate covalent interactions of compounds **1** and **4** with LC3/GABARAP as recently reported for **1** with the E3 ligase DCAF11^25^, we also studied the interaction of compound **1** and **4** with GABARAPL2 using ESI mass spectrometry, where no covalent modification was detected *in vitro* (data not shown).

Since compounds **1**-**4** were reported to trigger significant target degradation in cellular assays^6, 22, 26, 27^, we were interested in possible mechanisms causing these intriguing effects. In order to identify possible targets of these small molecules, we used an amine-linker adduct of the small molecules while using the same attachment point for linkers as in recently published ATTECs (Figure 3 D). For proteome-wide screening, we modified 10O5 (**4**) with a PEG-based linker and 8F20 (**3**) that both can be immobilized on sepharose beads to generate an affinity matrix for target pulldown, resulting in compounds **18** and **19**. As expected, dose-dependent competition assays showed that the KIF11 inhibitor 8F20/Ispinesib (**3**) selectively bound to KIF11 in HEK293T lysates (EC_50_ of 290 nM). No additional targets were detected for **19** confirming excellent selectivity of this inhibitor for KIF11 (Figure 3 D). Interestingly, GABARAPL2 was not significantly enriched in pull downs using **18** (up to 30 μM), suggesting that the interaction with this Atg8 homolog might be too weak to interact in cell lysates, confirming our hypothesis that the weak interaction detected by NMR was not sufficient to support a role as ATTEC based degrader. As additional cellular on target validation, we used the **3** and **4**-based tracer molecules **16** and **17** to measure cellular LC3A and LC3B target engagement using the NanoBRET technology, which again did not confirm binding of the 8F20 and 10O5 analogs to LC3A and LC3B in agreement with our assay data on recombinant proteins (SI Figures S7 A-C). To further study the cellular effects of the 4 ATTEC ligands (**1**-**4**), we monitored cellular growth in a live cell imaging system. Using RPE1 and U2OS cells, cell growth was monitored in live cells over a time course of 72 h after treatment with the respective compounds. Apart from **3**, no compound caused growth inhibition at concentrations < 10 μM, while compound **3** clearly suppressed cellular growth (SI Figures S7 D and E) without effecting cell viability at concentrations up to 30 μM (SI Figure S7 F). Since we have identified KIF11 as the only proteome-wide high affinity target and given the established role of this kinesin in cell division, KIF11 inhibition by **3** might trap cells in mitosis preventing cell division.^28^ Therefore, we analyzed the images taken during the live cell growth assay and found an increased number of rounded cells, indicating cells in a mitotic defect, consistent with the G2/M arrest as a result of 8F20 **(3)** treatment and in agreement with the literature (SI Figure S8).^28^

Intrigued by the proposed mechanism of action of ATTECs and the possible advantages over PROTACs (e.g. no complex ubiquitin transfer mechanism or higher hurdle for cells ATTEC resistance), we carried out screens for the identification of novel LC3/GABARAP binders, using two independent approaches. In the first approach, we initiated a virtual screening campaign by using a library of >7500 diverse in-house compounds. All the docking poses of *in silico* hits were individually inspected and we collected 281 compounds for experimental validation using the developed FP assay and all LC3/GABARAPs. Experimentally confirmed hits were studied using similarity search, which ultimately led to two novel LC3/GABARAP ligands (Figures 4 A and B). Interestingly, both ligands harbored two carboxylic acid moieties and initial SAR insights using 26 ligands of this compound class (SI Tables 1-3) revealed the importance of both carboxylic acids for binding. The first hit, LY223982 (**20**), has been developed targeting the leukotriene B4 receptor^29^ and showed selective binding to LC3 family members (Figures 4 B and C). Interestingly, TH152^30^ (**21**) displayed a K_D_ of 2 μM for LC3A in ITC titrations and interacted with all LC3/GABARAPs in FP assays (Figure 4 B and C). Therefore, TH152 represents the most potent reversible pan-LC3/GABARAP ligand reported so far.

**Figure 4:**
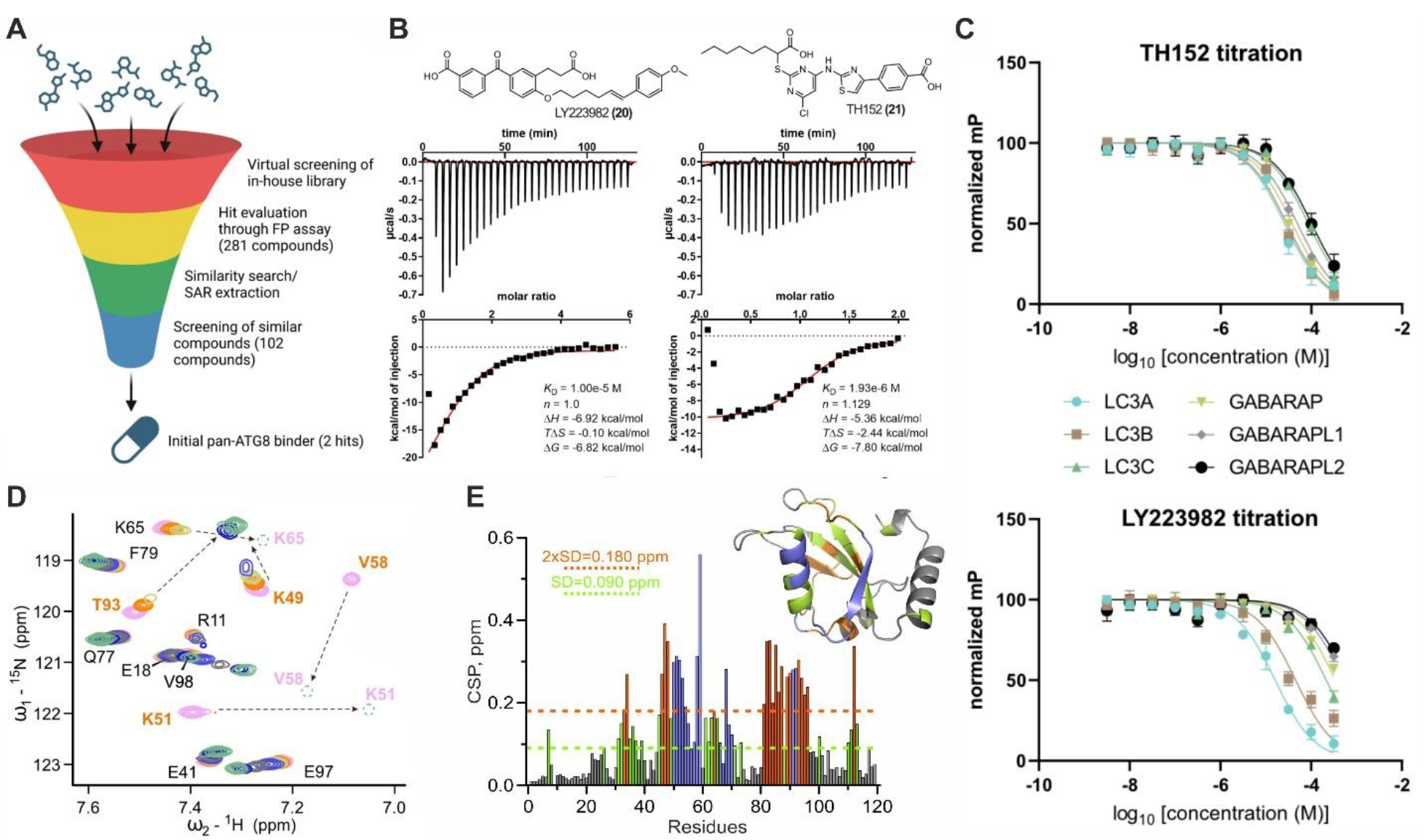
LC3/GABARAP hit identification campaigns via virtual screening. **A)** Schematic workflow of the virtual screening approach which was combined with biophysical hit validation. Our chemically diverse in-house library (>7500 compounds) was screened virtually using AutoDock^38^ and SeeSAR (BioSolveIT). 281 screening hits were screened against all LC3/GABARAPs using FP assay. Validated hits were used for similarity search within the in-house library was carried out using InfiniSee (BioSolveIT) and 102 similar compounds were screened again in vitro using FP assay resulting in two hits with affinity ≤ 10μM towards LC3A. **B)** Structural representation and corresponding ITC data for the two screening hits (compounds **20** and **21**). **C)** FP assay results using compounds **21** (upper panel) and **20** (lower panel) for selectivity screening within the human Atg8-family proteins. Data were measured in technical triplicates with error bars expressing the SD (n=3). **D)** Interaction between LC3B and compound **21** investigated by NMR. Representative areas of LC3B [^15^N,^1^H] BEST-TROSY spectra around the K51 and V58 backbone HN resonances are shown in overlay with free LC3B (magenta), and in stepwise increase of **21** molar ratios up to 1:4. Arrows show directions of large CSP for the resonances which are in the intermediate exchange mode. **E)** CSP values, induced by compounds **21** at molar ratio 1:2, are plotted against LC3B residue numbers. The light green dashed line indicates the standard deviations (SD) over all residues, the orange dashed line indicates double SD values. The blue bars are for LC3B residues which undergo strong intermediate exchange mode (significant decrease of the resonances intensity upon titration with **21**). Top right structure represents 3D mapping of CSP values on LC3B structure (pdb: 1UGM), indicating the HP2 as a most relevant interaction site. Residues with small (CSP < SD), intermediate (SD < CSP < 2xSD) or strong (2xSD < CSP) CSP values are marked in grey, light green and orange, respectively. LC3B residues which undergo strong intermediate exchange mode are marked blue. More details on this titration and titration of **21** to the GABARAP protein are depicted in SI Figure S9.

To validate pan-Atg8 binding activity, we characterized the interaction of TH152 with ^15^N labelled LC3B and GABARAP protein by NMR titrations (as exemplary depicted in Figure 4 D-F for LC3B with full NMR data analysis shown in SI Figure S9). The NMR results directly indicate that LC3B and GABARAP interact with TH152 over LDS, confirming molecular modelling of these interactions. However, due to the presence of two carboxylic acid groups the identified ligands might be characterized by poor cell penetration and might therefore require optimization for ATTEC development.

As a second hit finding approach, we conducted a large-scale fragment screening campaign using X-ray crystallography by soaking a total of 1006 LC3B crystals with a diverse fragment library. This led to the collection of over 800 high quality diffraction datasets, which identified a total of 21 diverse hits in the binding cavities on LC3B after refinement of the structures (Figure 5A). This set significantly complements earlier fragment and hit finding campaigns using NMR and DEL (DNA encoded library) screening.^20^ Our screen confirmed that HP2 is the most druggable binding site on LC3/GABARAP proteins surface, accommodating 10 from 21 identified fragments (Figure 5B). Fragments such as x0145 (HP1) and x0626 (S2) offer the possibility for fragment linking. Previously reported data^20^ displays three fragments bound to HP2 with one fragment bound twice as depicted in Figure 5C. Interestingly, all of the previously published fragments also harbor a carboxylic acid group. Comparison with hit rates of similar protein interaction domains such as E3 ligases suggest that the HP2 pocket can accommodate a diversity of ligands, indicating a good druggability of this site. Based on our success with the very limited *in silico* study and the fragment screening campaign, we concluded that design and development of potent LC3/GABARAP ligands for ATTECs should be feasible. Making this rich pool of hit matter available together with the established assay platform will allow robust validation of LC3/GABARAP ligands which may be developed to either selectively target one human Atg8 family member or to design pan-Atg8 ligands.

**Figure 5:**
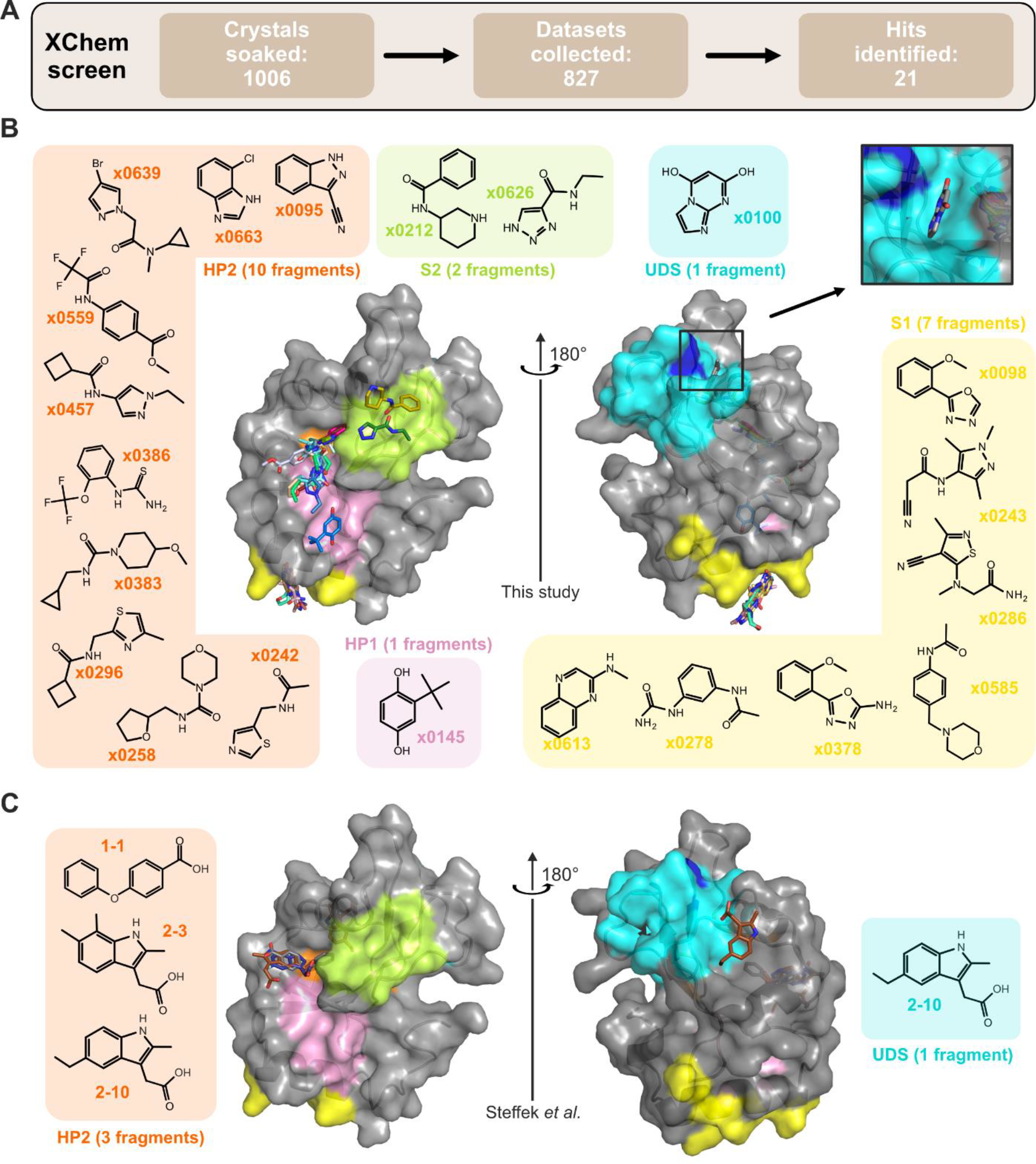
LC3/GABARAP hit identification via X-ray crystallography fragment screening (XChem). **A)** Schematic workflow of the XChem screening, resulting in 21 identified hits. **B)** Overlay of crystal structures containing diverse fragments bound to LC3B. Bound fragments are depicted as chemical structures with colored background corresponding to the binding site: HP1 (pink), HP2 (orange), UDS (cyan, key UDS residue F80 is shown blue), new identified regions S1 (yellow) and S2 (light green). The insert shows the correct pose for x0100 within UDS (rotations by y45° and x20° degrees from the main plot). All structures available at protein data bank (PDB IDs 7GA8-7GA9 and 7GAA-7GAS). Exemplary electron density maps for representative binders are shown in SI Figure S10. **C)** Overlay of the three published crystal structures of LC3A containing small molecule fragments (pdb: 7R9W, 7R9Z and 7RA0).^20^

## Discussion

In this study we validated published LC3/GABARAP ligands that could serve as starting points for the development of more potent ligands as well as ATTECs. To our surprise, none of the ligands used for the development of ATTECs interacted with LC3/GABARAP, suggesting that reported degraders did not function as LC3/GABARAP modulators causing degradation by the autophagy/lysosomal pathway. The hypothesis using LC3/GABARAP as receptors for degrader development is quite new. As a result, few tools are available that could be utilized demonstrating that observed degradation events indeed are associated with autophagy. In the ubiquitin based degrader field, pathway association of PROTACs is usually accomplished using a) inactive E3 ligands such as the inactive stereoisomer of VHL or N-methylated thalidomide derivatives.^31^ These tools account also for possible effect caused by POI ligand that is often a pharmacological highly active small molecule; b) proteasomal inhibitors that rescue proteasome dependent degradation events; c) inhibitors of E3 ligase activating enzymes such as neddylation inhibitors for cullin dependent E3 ligases^32^ and finally a proteome wide analysis demonstrating degrader selectivity. However, for ATTECs, pathway activators such as mTOR inhibitors or small molecules such as Bafilomycin could be used. We strongly believe that for new ligands common community guidelines for chemical probe developments should be applied. These quality standard would comprise direct on-target engagement assays and for stronger ligands also for cell based assay systems, appropriate controls comprising inactive control molecules, drug resistant mutants or knock out cell lines.^33-36^ Recently also first standards for covalent inhibitors and degrader molecules have been defined.^37^

Our druggability analysis, *in silico* and fragment screening revealed that LC3/GABARAP are druggable and first low μM ligands that were confirmed in our assays have been reported. In our comprehensive fragment screening study, diverse ligands binding to the HP2 site emerged, but also HP1 and initial ligands for the UIM site were identified. It is our hope that fragment growing and linking efforts will result in more potent LC3/GABARAP ligands that could be used for the development of efficient ATTECTs in the future.

## Author contributions

Manuscript and figures were prepared by MPS and edited by VVR and SK. MPS, JD, AK, FAG, JH, SC, FL, NB, SL, IB, VM, DF, PGM, CWET and LB contributed experimental data. Scientific supervision by TH, NA, FvD, AS, BK, SMK, KS, SK, EP and VVR.

## Acknowledgements

Compounds 1-4 were kindly provided by Akinori Toita and Misaki Homma (Takeda). MPS, AK, FAG, NB, TH, VM, JD, KS, SMK, SK and VVR are grateful for support by the Structural Genomics Consortium (SGC), a registered charity (no: 1097737) that receives funds from Bayer AG, Boehringer Ingelheim, Bristol Myers Squibb, Genentech, Genome Canada through Ontario Genomics Institute, EU/EFPIA/OICR/McGill/KTH/Diamond Innovative Medicines Initiative 2 Joint Undertaking [EUbOPEN grant 875510], Janssen, Merck KGaA, Pfizer and Takeda and by the German Cancer Research Center DKTK and the Frankfurt Cancer Institute (FCI). MPS is funded by the Deutsche Forschungsgemeinschaft (DFG, German Research Foundation), CRC1430 (Project-ID 424228829). SL and BK are funded by the Deutsche Forschungsgemeinschaft (DFG) SFB1309 (ID: 325871075, project ID: 401883058). The authors would like to acknowledge the Diamond Light Source for access to the fragment screening facility XChem, for usage of DSi-Poised library and for beam time on beamline I04-1 under proposal LB29658. AS, SK and SC are grateful for support by the BMBF program PROXIDRUGs. Figures were prepared using BioRender.

## Methods

### Chemistry

#### (*R*)-*N*-(1-amino-12-oxo-3,6,9-trioxa-13-azahexadecan-16-yl)-*N*-(1-(3-benzyl-7-chloro-4-oxo-3,4-dihydroquinazolin-2-yl)-2-methylpropyl)-4-methylbenzamide (compound 19)

A mixture of (*R*)-*N*-(3-aminopropyl)-*N*-(1-(3-benzyl-7-chloro-4-oxo-3,4-dihydroquinazolin-2-yl)-2-methylpropyl)-4-methylbenzamide (Ispinesib, 80 mg, 150 μmol), (7-Azabenzotriazol-1-yloxy)tripyrrolidinophosphonium hexafluorophosphate (123 mg, 216 μmol), 2,2-dimethyl-4-oxo-3,8,11,14-tetraoxa-5-azaheptadecan-17-oic acid (52 mg, 162 μmol) and *N,N*-diisopropylethylamine (40 μL, 232 μmol) in anh. DMF (5 mL) was stirred at ambient temperature. After 1 h, the mixture was partitioned between water and ethyl acetate. The ethyl acetate layer was washed with brine, dried over MgSO_4_, filtered, and volatiles were removed under reduced pressure. The residue was dissolved in DCM/TFA (3/1, 8 mL) and stirred for 1 h. After 1 h, all volatiles were removed under reduced pressure to provide the title compound (100 mg, 90%) which was used in the next step without further purification. MS (ESI): m/z calc. for [M+H^+^]^+^ = 720.34, found = 720.30.

#### (*R*)-*N*-(1-(3-benzyl-7-chloro-4-oxo-3,4-dihydroquinazolin-2-yl)-2-methylpropyl)-*N*-(1-(5,5-difluoro-7-(1*H*-pyrrol-2-yl)-5*H*-5l4,6l4-dipyrrolo[1,2-c:2’,1’-f][1,3,2]diazaborinin-3-yl)-3,16-dioxo-7,10,13-trioxa-4,17-diazaicosan-20-yl)-4-methylbenzamide (compound 17)

A mixture of (*R*)-*N*-(1-amino-12-oxo-3,6,9-trioxa-13-azahexadecan-16-yl)-*N*-(1-(3-benzyl-7-chloro-4-oxo-3,4-dihydroquinazolin-2-yl)-2-methylpropyl)-4-methylbenzamide (15 mg, 21 μM), 2,5-dioxopyrrolidin-1-yl 3-(5,5-difluoro-7-(1*H*-pyrrol-2-yl)-5*H*-5λ^4^,6λ^4^-dipyrrolo[1,2-*c*:2’,1’-*f*][1,3,2]diazaborinin-3-yl)propanoate (8.5 mg, 20 μM) and *N,N*-diisopropylethylamine (8.6 μL, 50 μM) in anh. DMF (0.3 mL) was stirred at ambient temperature for 2 h and afterwards purified by prep. HPLC (H_2_O/ACN with 0.1% TFA) to provide the title compound (18 mg, 88%). ^1^H NMR (500 MHz, DMSO-*d*_6_): δ 11.50 (s, 1H), 8.23 (d, *J* = 8.6 Hz, 1H), 7.81 (d, *J* = 2.1 Hz, 1H), 7.79 (s, 1H), 7.66 (dd, *J* = 8.6, 2.1 Hz, 1H), 7.46 (s, 1H), 7.44 – 7.39 (m, 2H), 7.38 (d, *J* = 4.6 Hz, 2H), 7.36 (d, *J* = 7.6 Hz, 2H), 7.34 – 7.27 (m, 2H), 7.28 – 7.19 (m, 7H), 7.01 (d, *J* = 4.0 Hz, 1H), 6.45 (d, *J* = 3.9 Hz, 1H), 6.37 (dt, *J* = 4.2, 2.2 Hz, 1H), 5.88 (d, *J* = 16.2 Hz, 1H), 5.54 (d, *J* = 10.5 Hz, 1H), 5.05 (d, *J* = 16.3 Hz, 1H), 3.57 (t, *J* = 5.4 Hz, 2H), 3.56 – 3.50 (m, 3H), 3.49 – 3.39 (m, 5H), 3.27 (q, *J* = 10.5, 8.9 Hz, 5H), 3.15 (t, *J* = 7.5 Hz, 3H), 2.96 (q, *J* = 5.5 Hz, 2H), 2.73 (dq, *J* = 10.8, 6.4 Hz, 1H), 2.33 (s, 3H), 2.03 (q, *J* = 6.8 Hz, 2H), 1.35 – 1.14 (m, 2H), 0.89 (d, *J* = 6.7 Hz, 3H), 0.87 – 0.78 (m, 1H), 0.47 (d, *J* = 6.3 Hz, 3H). ^13^C NMR (126 MHz, DMSO): δ 171.99, 170.21, 169.48, 168.30, 161.13, 155.24, 152.40, 150.96, 147.21, 139.52, 138.71, 137.44, 136.69, 133.77, 133.08, 133.01, 128.91, 128.71, 128.67, 128.04, 127.45, 126.71, 126.42, 126.13, 125.90, 124.41, 122.78, 119.96, 119.10, 118.07, 118.02, 117.96, 116.07, 111.80, 69.67, 69.64, 69.60, 69.44, 66.67, 66.64, 58.99, 45.18, 42.32, 35.83, 35.77, 29.33, 25.47, 22.99, 20.89, 19.48, 18.16. HRMS (MALDI): m/z calc. for [M+Na^+^]^+^ = 1053.4383, found = 1053.4377

#### *Tert*-butyl 2-(2,6-dibromo-4-formylphenoxy)acetate

3,5-dibromo-4-hydroxybenzaldehyde (5.6 g, 20 mmol) was solved in anh. DMF (60 mL). K_2_CO_3_ (5.52 g, 40 mmol) was added and the mixture was stirred for 5 min. *Tert*-butyl 2-bromoacetate (4,68g, 24 mmol) was stirred at ambient temperature overnight until complete consumption of starting material. The mixture was partitioned between water and ethyl acetate. The ethyl acetate layer was washed with brine, dried over MgSO_4_, filtered, and volatiles were removed under reduced pressure yielding a pale yellow oil which crystallized overnight. The product was used without further purification (92%) MS (ESI): m/z calc. for [C_13_H_14_Br_2_O_4_ +Na^+^]^+^ = 417.06, found = 416.85 ^1^H NMR (500 MHz, DMSO-*d*_*6*_) δ 9.90 (s, 1H), 8.17 (s, 2H), 4.64 (s, 2H), 1.46 (s, 9H). ^13^C NMR (126 MHz, DMSO-*d*_*6*_) δ 190.04, 190.02, 166.06, 156.20, 134.57, 133.73, 118.19, 81.86, 69.42, 27.67.

#### *Tert*-butyl 2-(2,6-dibromo-4-((5-iodo-2-oxoindolin-3-ylidene)methyl)phenoxy)acetate

A mixture of *tert*-butyl 2-(2,6-dibromo-4-formylphenoxy)acetate (0.606 g, 1.5 mmol), 5-iodoindolin-2-one (0.518 g, 2 mmol) were suspended in absolute ethanol (8 mL). Catalytic amounts of piperidine (0.1 eq, 0.013 g, 0,15 mmol or 15 μL) were added and the mixture was refluxed at 80°C for 3h. After 3h an orange solid formed. The solid was filtered through a glass frit and rinsed with cold ethanol. The solid was collected and used in the next step without purification yielding an inseparable mixture of *E/Z* isomers. The mother liquor was evaporated *in vacuo* and purified by flash chromatography (*n*-hexanes/EtOAc, 6:1) to increase the overall yield (80%). MS (ESI): m/z calc. for [C_13_H_14_Br_2_O_4_ +Na^+^]^+^ = 658,09, found = 657.80. ^1^H NMR (400 MHz, DMSO-*d6*) δ 10.77 (s, 1H), 8.78 (s, 1H), 8.01 (d, J = 0.7 Hz, 2H), 7.72 (d, J = 1.7 Hz, 1H), 7.60 – 7.52 (m, 3H), 6.73 (d, J = 8.2 Hz, 1H), 4.62 (d, J = 0.7 Hz, 2H), 4.59 (d, J = 0.7 Hz, 1H), 1.47 (dd, J = 2.2, 0.7 Hz, 13H). ^13^C NMR (101 MHz, DMSO-*d6*) δ 166.49, 166.23, 152.93, 140.54, 138.65, 137.53, 136.19, 134.44, 133.58, 133.53, 133.38, 133.08, 130.65, 128.44, 128.03, 127.09, 126.96, 117.51, 116.84, 112.71, 111.98, 84.20, 81.78, 69.50, 69.48, 27.74, 27.72.

#### 2-(2,6-dibromo-4-((5-iodo-2-oxoindolin-3-ylidene)methyl)phenoxy)acetic acid

*Tert*-butyl 2-(2.6-dibromo-4-formylphenoxy)acetate (0.300 mg, 0.47 mmol) was solved in absolute DCM (10 mL). TFA (2 mL) was added dropwise to the solution and let stir at ambient temperature for 2 h until complete consumption of starting material. After 2 h a red solid formed which was transferred into a glass frit and rinsed witch cold DCM. The crystals were collected and dried *in vacuo* overnight (95%). ^1^H NMR (400 MHz, DMSO-*d6*) δ 13.17 (s, 1H), 10.83 (s, 1H), 8.78 (s, 2H), 8.06 – 7.99 (m, 2H), 7.85 (s, 1H), 7.71 (d, J = 1.7 Hz, 1H), 7.61 – 7.51 (m, 2H), 6.71 (dd, J = 20.1, 8.1 Hz, 2H), 4.62 (s, 1H), 4.60 (s, 2H). ^13^C NMR (101 MHz, DMSO-*d6*) δ 168.56, 167.58, 166.49, 152.85, 142.79, 140.54, 138.64, 137.54, 136.18, 134.45, 133.60, 133.52, 133.14, 130.70, 128.45, 128.08, 127.09, 126.97, 123.04, 117.66, 116.96, 112.70, 111.98, 84.21, 83.87, 68.86, 68.83.

#### *Tert*-butyl(1-(2,6-dibromo-4-((5-iodo-2-oxoindolin-3-ylidene)methyl)phenoxy)-2-oxo-7,10,13-trioxa-3-azahexadecan-16-yl)carbamate

A mixture of 2-(2,6-dibromo-4-((5-iodo-2-oxoindolin-3-ylidene)methyl)phenoxy)acetic acid (0.316 g, 0.55 mmol), *tert*-butyl (3-(2-(2-(3-aminopropoxy)ethoxy)ethoxy)propyl)carbamate (0.192 g, 0.6 mmol) were solved in anhy. DMF (18 mL). PyAOP (0.342 g, 0.65 mmol) and DIPEA (0,091 g, 0,71 mmol or 125 μL) were added to the mixture and let stir for 1.5h at ambient temperature. The crude mixture was evaporated *in vacuo* and purified directly via reverse phase column chromatography (H_2_O/ACN) (79%). MS (ESI): m/z calc. for [C_13_H_14_Br_2_O_4_ +Na^+^]^+^ = 904,4 found = 904.00 ^1^H NMR (500 MHz, Methylene Chloride-*d2*) δ 9.27 (s, 1H), 8.51 (s, 1H), 7.85 – 7.73 (m, 2H), 7.55 – 7.47 (m, 1H), 7.39 – 7.31 (m, 1H), 7.25 (s, 1H), 6.69 (dd, J = 25.4, 8.2 Hz, 1H), 5.17 – 4.97 (m, 1H), 4.56 (s, 1H), 4.52 (s, 1H), 3.60 (d, J = 5.9 Hz, 6H), 3.57 – 3.43 (m, 9H), 3.16 (q, J = 6.4 Hz, 2H), 1.88 (td, J = 6.3, 3.3 Hz, 2H), 1.70 (p, J = 6.2 Hz, 2H). ^13^C NMR (126 MHz, Methylene Chloride-*d2*) δ 167.54, 167.47, 156.48, 153.39 (d, J = 39.1 Hz), 140.71, 139.54, 138.62, 136.74, 134.33, 134.14, 134.01, 132.22, 128.97, 118.70, 117.82, 113.11, 112.51, 79.13, 71.81, 71.73, 71.04, 71.02, 71.01, 70.99, 70.96, 70.71, 70.67, 70.07, 70.01, 69.93, 69.89, 54.43, 54.22, 54.00, 53.78, 53.57, 39.04, 37.77, 37.74, 30.31, 29.82, 29.79, 28.74.

#### N-(3-(2-(2-(3-aminopropoxy)ethoxy)ethoxy)propyl)-2-(2,6-dibromo-4-((5-iodo-2-oxoindolin-3-ylidene)methyl)phenoxy)acetamide (TFA salt) (compound 18)

*Tert*-butyl (1-(2,6-dibromo-4-((5-iodo-2-oxoindolin-3-ylidene)methyl)phenoxy)-2-oxo-7,10,13-trioxa-3-azahexadecan-16-yl)carbamate (0.021 g, 0.024 mmol) was charged into a flask and solved in anhy. DCM (1 mL). TFA (0.6 mL) was added and the solution for stirred for 1h until complete consumption of starting material. Toluene (2 mL) was added and the solution was evaporated and dried *in vacuo* overnight yielding the title compound as TFA salt. MS (ESI): m/z calc. for [C_43_H_44_BBr_2_F_2_IN_6_O_7_+H^+^]^+^= 782.28 found = 782.05

#### *N*-(1-(2,6-dibromo-4-((5-iodo-2-oxoindolin-3-ylidene)methyl)phenoxy)-2-oxo-7,10,13-trioxa-3-azahexadecan-16-yl)-3-(5,5-difluoro-7-(1H-pyrrol-2-yl)-5H-4l4,5l4-dipyrrolo[1,2-c:2’,1’-f][1,3,2]diazaborinin-3-yl)propenamide (compound 16)

*Tert*-butyl (1-(2,6-dibromo-4-((5-iodo-2-oxoindolin-3-ylidene)methyl)phenoxy)-2-oxo-7,10,13-trioxa-3-azahexadecan-16-yl)carbamate (0.025 g, 0.029 mmol) was charged into a flask and solved in anhy. DCM (1 mL). TFA (0.7 mL) was added and the solution for stirred for 1h until complete consumption of starting material. Toluene (2 mL) was added and the solution was evaporated *in vacuo* and used directly in the next step without purification. The crude was solved in anhy. DMF (1 mL) and the flask was wrapped in tin foil. Py-BODIPY-NHS ester (0.011 g, 0.026 mmol) and DIPEA (0.06 mM,11 μL) were added to the solution and stirred for 2 h at ambient temperature. The crude was afterwards purified by prep. HPLC (H_2_O/ACN with 0.1% TFA) to provide the title compound (80%). MS (HRMS): m/z calc. for [C_43_H_44_BBr_2_F_2_IN_6_O_7_+Na^+^]^+^= 1113,0744, found = 1113,06312. ^1^H NMR (400 MHz, DMSO-*d6*) δ 11.40 (s, 1H), 10.84 (s, 1H), 8.79 (s, 2H), 8.17 (dt, J = 11.8, 5.8 Hz, 2H), 8.03 (dd, J = 4.9, 1.2 Hz, 2H), 7.90 (t, J = 5.6 Hz, 1H), 7.86 (s, 1H), 7.57 (td, J = 8.2, 1.7 Hz, 2H), 7.43 (s, 1H), 7.38 – 7.32 (m, 3H), 7.27 (td, J = 2.7, 1.3 Hz, 1H), 7.16 (d, J = 4.5 Hz, 1H), 7.01 (d, J = 4.0 Hz, 1H), 6.69 (d, J = 8.1 Hz, 1H), 6.37 – 6.29 (m, 3H), 4.45 (s, 1H), 4.43 (s, 2H), 3.55 – 3.43 (m, 16H), 3.25 (q, J = 6.7, 5.0 Hz, 5H), 3.11 (dd, J = 12.9, 6.9 Hz, 6H), 1.73 (td, J = 6.7, 3.2 Hz, 3H), 1.64 (q, J = 6.6 Hz, 3H), 1.25 (d, J = 6.8 Hz, 4H). ^13^C NMR (101 MHz, DMSO-*d6*) δ 171.29, 166.97, 166.64, 156.43, 153.24, 150.69, 141.04, 139.12, 138.06, 137.41, 136.65, 134.87, 133.98, 133.76, 133.48, 132.87, 128.94, 127.55, 127.22, 126.59, 124.86, 123.37, 119.82, 117.90, 117.53, 116.60, 112.48, 111.99, 84.70, 71.46, 70.25, 70.10, 70.00, 68.86, 68.55, 36.56, 36.34, 34.39, 29.81, 29.64, 29.49, 29.18, 24.54, 22.56.

##### Protein expression and purification for biophysical assays

LC3A_1-120_, LC3B_1-120_, LC3C_1-126_, GABARAP_1-116_, GABARAPL1_1-117_ and GABARAPL2_1-117_ were expressed as a recombinant fusion protein incorporating a His6 and TEV cleavage site at the N-terminus. *E. coli* Rosetta cells were cultured in Terrific Broth (TB) at 37 °C until an OD600 of 1.0 was reached. The culture was then cooled to 18 °C and allowed to reach an OD600 of 2.5. Protein expression was induced by the addition of 0.5 mM isopropyl β-D-1-thiogalactopyranoside (IPTG) and the protein was allowed to express overnight. Cells were harvested (Beckman centrifuge, via centrifugation at 6000 g at 4°C) and lysed by sonication (SONICS vibra cell, 5 s on-, 10 s off cycle using a total of 30 minutes) in the presence of DNase I (Roche, Basel, CH) and cOmplete EDTA-free protease inhibitor (Roche, Basel, CH), and recombinant protein was purified using Ni-NTA-affinity chromatography in Purification buffer (30 mM 4-(2-hydroxyethyl)-1-piperazineethanesulfonic acid; pH 7.5 (HEPES), 500 mM NaCl, 5 % glycerol, 0.5 mM tris(2-carboxyethyl)phosphine (TCEP) and 30 mM Imidazole) and elution was carried out using Purification buffer including additional 300 mM Imidazole. The eluted proteins were dialyzed overnight into gel filtration buffer (30 mM HEPES pH 7.5, 250 mM NaCl, 5 % glycerol and 0.5 mM TCEP) while the expression tag was cleaved using 1 mg tobacco etch virus (TEV) protease. The cleaved protein was passed through a HiLoad® 26/600 Superdex® 75 pg (GE Healthcare) size exclusion chromatography column and the resulting pure protein was stored in gel filtration buffer, flash frozen in liquid nitrogen and subsequently stored at −80 °C for further experiments.

##### Protein purification for X-ray crystallography

Human LC3B (1-120) was cloned into the pNIC28-Bsa4 vector using restriction sites Lic5 and Lic3 to add a TEV-cleavable N-terminal His-tag. The construct was then transformed into E. coli Rosetta(DE3) competent cells and expressed in TB medium by overnight induction with 0.2 mM IPTG (OD600=2.5). Cells were harvested by centrifugation, resuspended in buffer A (30 mM HEPES pH 7.5 @ 4°C, 500 mM NaCl, 0.5 mM TCEP, 10 mM Imidazole, and 5% Glycerol), and lysed by sonication on ice. The soluble fraction was collected by centrifugation at 21000 g for 40 min. The fraction was with 4 ml Ni-NTA beads (pre-equilibrated with lysis buffer) for batch binding on ice for 1 hour. The beads were washed with buffer B (30 mM HEPES pH 7.5 @ 4°C, 500 mM NaCl, 0.5 mM TCEP, 30 mM Imidazole, and 5% Glycerol) and eluted with buffer C (30 mM HEPES pH 7.5 @ 4°C, 500 mM NaCl, 0.5 mM TCEP, 300 mM Imidazole, and 5% Glycerol). Protein in the eluted fraction was treated with TEV protease overnight while dialyzing against (30 mM HEPES pH 7.5 @ 4°C, 300 mM NaCl, 0.5 mM TCEP, and 5% Glycerol) to cleave the His-tag. The dialyzed mixture was passed through 4 ml Ni-NTA beads, flowthrough was collected, concentrated, and injected into GE Superdex 75 16/600 Prep grade column pre-equilibrated with SEC buffer (30 mM HEPES pH 7.5 @ 4°C, 100 mM NaCl, 0.5 mM TCEP, and 5% Glycerol). The peak was collected and concentrated to 22.5 mg/ml.

##### Isothermal Titration Calorimetry (ITC)

ITC experiments were performed using a NanoITC instrument (TA Instruments, New Castle, USA) at 25 °C in gel filtration buffer (50 mM Na2HPO4 pH = 7.0, 100 mM NaCl and 0.5 mM TCEP). 25 μM inhibitor dissolved in gel filtration buffer was titrated into purified LC3/GABARAPs at a concentration of 500 μM in the reaction cell. For this protocol, the chamber was pre-equilibrated with the protein, and the test compounds were titrated in while continuously measuring the rate of exothermic heat evolution. The heat of binding was integrated, corrected, and fitted to an independent single-binding site model based on the manufacturer’s instructions, from which thermodynamic parameters (ΔH and TΔS), equilibrium association and dissociation constants (K_A_ and K_D_, respectively), and stoichiometry (n) were calculated. Data were displayed using GraphPad Prism 9.3.

##### Differential scanning fluorimetry (DSF) assay

Differences in the melting temperature (ΔTm) data were measured as described in Schwalm *et al*.^39^ Purified proteins were buffered in DSF buffer (25 mM HEPES pH 7.5, 500 mM NaCl) and were assayed in a 384-well plate (Thermo, #BC3384) with a final protein concentration of 20 μM in 10 μL final assay volume. Inhibitors were added in excess to a final concentration of 40 μM, using an ECHO 550 acoustic dispenser (Labcyte). As a fluorescent probe, SYPRO-Orange (Molecular Probes) was used at 5x final concentration. Filters for excitation and emission were set to 465 nm and 590 nm, respectively. The temperature was increased from 25 °C with 3 °C/min to a final temperature of 99 °C, while scanning, using the QuantStudio5 (Applied Biosystems). Data was analyzed using Boltzmann-equation in the Protein Thermal Shift software (Applied Biosystems). Samples were measured in technical triplicates.

#### Affinity determination using spectral shift mode on Dianthus

Protein Labeling Kit RED-maleimide 2nd Generation (cat# MO-L014; NanoTemper Technologies GmbH) was used for covalent labeling of LC3A cystine residues. Labelling was carried out following the manufacturer protocol, using a 3:1 ratio dye:protein. Labeled LC3A with degree of labeling of 0.7 (as determined by UV-VIS absorbance spectroscopy) was purified in 30 mM HEPES, 100mM NaCl, 0.05% Tween, pH 7.5. For spectral shift assays, a 16-point affinity measurement was performed in duplicates using the Dianthus (NanoTemper Technologies GmbH, Germany) instrument. Assays were performed in 10 mM Na2HPO4/KH2PO4, 0.05% Tween-20, 5 % DMSO, pH 7.4 with a maximum ligand concentration of of 500 μM. All compounds dissolved DMSO were first pre-diluted to 1 mM in assay buffer followed by a 16-fold 1:1 dilution series of each compound in a Dianthus 384 micro plate. Measurements were performed in spectral shift mode with an LED excitation power of 100%. Data were analyzed using the DI.Screening Analysis Software (v.2.0.4) (NanoTemper Technologies GmbH, Germany) and quality criteria values including, Δ Ratio, Signal-to-Noise-Ratio, Saturation and Kd values were determined.

##### Fluorescence polarization assay (FP assay)

For the complementation assay, the fluorescently labeled p62 LIR probe (SDNSSGGDDDWTHLSSK-Cy5) was diluted to (30 nM) in assay buffer (50 mM HEPES pH 7.5, 150 mM NaCl, 5 % glycerol, 1 mM TCEP and 0.05% TWEEN20) in a black 384-well flat bottom plate (Greiner Bio-One, #784076) and purified LC3/GABARAPs were titrated in a concentration range from 55 μM to 600 pM. After 1 h incubation at room temperature, fluorescence polarization was measured with polarized excitation wavelength of 590 nm and filtered emission wavelength of 675 nm, respectively, using a PHERAstar plate reader (BMG Labtech). Resulting data was plotted using GraphPad Prism 9.3 software and analyzed using a nonlinear fit to calculate the probe IC_50_. For competition assays, 30 nM probe was added to assay buffer containing 4 μM LC3/GABARAPs. Compounds were titrated from 20 μM to 20 nM using an ECHO 550 acoustic dispenser (Labcyte) incubated for 1 h at room temperature and subsequent read out as described above. Data was plotted in GraphPad Prism 9.3 and analyzed using a nonlinear fit (equation: Y=100/(1+10^(X-LogIC50)) for IC50 determination. KI calculation was performed using the Nikolovska-Coleska formula.^40^

##### Covalent compound screening

For screening of covalent compounds, 46 compounds were tested against all LC3/GABARAPs, purified as described above. For LC-MS experiments, 50 μM of protein was used together with 100 μM of compound. The reaction was incubated for 90 min at room temperature and stopped by a 1:30 dilution in H_2_O with 0.1 % formic acid. Samples were measured, using an Agilent 6230 TOF LC/MS. Data was evaluated using the BioConfirm B.08.00 software.

##### NanoBRET cellular target engagement assay

The assay was performed as described previously.^41^ In brief: Constructs contained the cDNA of full-length LC3A and LC3B cloned in frame with an N-terminal NanoLuc-fusion. Plasmids were transfected into HEK293T cells using FuGENE HD (Promega, E2312) and proteins were allowed to express for 20 h. 10O5 and 8F20-based tracers were titrated to the protein as depicted in SI Figure 4 D. For competition experiments, 1 μM of the tracers was pipetted into white 384-well plates (Greiner 781 207) using an Echo 550 acoustic dispenser (Labcyte) containing LC3A/LC3B expressing transfected cells at a density of 2.5x10^5^ cells/mL in Opti-MEM without phenol red (Life Technologies). The system was allowed to equilibrate for 2 hours at 37 °C and 5% CO_2_ prior to BRET measurements. To measure BRET, NanoBRET NanoGlo Substrate + Extracellular NanoLuc Inhibitor (Promega, N2540) was added as per the manufacturer’s protocol, and filtered luminescence was measured on a PHERAstar plate reader (BMG Labtech) equipped with a luminescence filter pair (450 nm BP filter (donor) and 610 nm LP filter (acceptor)). Competitive displacement data were then plotted using GraphPad Prism 9.3 software using a normalized 3-parameter curve fit with the following equation: Y=100/(1+10^(X-LogIC50)).

##### Preparation of Affinity Matrix 10O5 and 8F20 (5)

Compounds **18** and **19** (1 μM) were linked to DMSO-washed NHS-activated (∼20 μM/mL beads) sepharose beads (1 mL) and triethylamine (20 μL) in DMSO (2 mL) on an end-over-end shaker overnight at RT in the dark. Aminoethanol (50 μL) was then added to inactivate the remaining NHS-activated carboxylic acid groups. After 16 hours the beads were washed with 10 mL DMSO and 30 mL EtOH to yield an affinity matrix of 10O5 and 8F20, respectively which were stored at 4 °C in EtOH. Successful immobilization was controlled by LC-MS and Kaiser-test.^42^

##### Preparation of cell lysates for affinity pulldown assays

HEK293 cells were cultured in DMEM (PAN Biotech). All media were supplemented in with 10% FBS (PAN Biotech) and cells were internally tested for Mycoplasma contamination. Cells were lysed in lysis buffer (0.8% Igepal, 50 mM Tris-HCl pH 7.5, 5% glycerol, 1.5 mM MgCl_2_, 150 mM NaCl, 1 mM Na_3_VO_4_, 25 mM NaF, 1 mM DTT and supplemented with protease inhibitors (SigmaFast, Sigma) and phosphatase inhibitors (prepared in-house according to Phosphatase inhibitor cocktail 1, 2 and 3 from Sigma-Aldrich)). The protein amount of cell lysates was determined by Bradford assay and adjusted to a concentration of 5 mg/mL.^42^

##### Competition pulldown assays

For the selectivity profiling of free Compound **18** and **19**, lysates from HEK293 cells were adjusted to 5 mg/mL protein concentration (0.4% Igepal). Then, 0.5 mL lysate was pre-incubated with 10 doses of the compounds (DMSO vehicle, 3 nM, 10 nM, 30 nM, 100 nM, 300 nM, 1000 nM, 3000 nM, 10000 nM, 30000 nM) for 1 h at 4°C in an end-over-end shaker, followed by incubation with 18 μL the Affinity Matrix 10O5 and 8F20 for 30 min at 4 °C in an end-over-end shaker. ^42^

The beads were washed (1x 1 mL of lysis buffer without inhibitors and only 0.4% Igepal, 2x 2mL of lysis buffer without inhibitors and only 0.2% Igepal) and captured proteins were denatured with 8 M urea buffer, alkylated with 55 mM chloroacetamide and digested with Trypsin according to standard procedures. Resulting peptides were desalted on a C18 filter plate (Sep-Pak® tC18 μElution Plate, Waters), vacuum dried and stored at -20 °C until LC-MS/MS measurement.

##### LC-MSMS measurement of (competition) pulldown assays

Peptides were analyzed via LC-MS/MS on a Dionex Ultimate3000 nano HPLC coupled to an Orbitrap Fusion Lumos mass spectrometer, operated via the Thermo Scientific Xcalibur software. Peptides were loaded on a trap column (100 μm x 2 cm, packed in house with Reprosil-Gold C18 ODS-3 5 μm resin, Dr. Maisch, Ammerbuch) and washed with 5 μL/min solvent A (0.1 % formic acid in HPLC grade water) for 10 min. Peptides were then separated on an analytical column (75 μm x 40 cm, packed in house with Reprosil-Gold C18 3 μm resin, Dr. Maisch, Ammerbuch) using a 50 min gradient ranging from 4-32 % solvent B (0.1 % formic acid, 5 % DMSO in acetonitirile) in solvent A (0.1 % formic acid, 5 % DMSO in HPLC grade water) at a flow rate of 300 nL/min.

The mass spectrometer was operated in data dependent mode, automatically switching between MS1 and MS2 spectra. MS1 spectra were acquired over a mass-to-charge (m/z) range of 360-1300 m/z at a resolution of 60,000 (at m/z 200) in the Orbitrap using a maximum injection time 50 ms and an automatic gain control (AGC) target value of 4e5. Up to 12 peptide precursors were isolated (isolation width of 1.2 Th, maximum injection time of 75 ms, AGC value of 2e5), fragmented by HCD using 25 % 30% normalized collision energy (NCE) and analyzed in the Orbitrap at a resolution of 15,000. The dynamic exclusion duration of fragmented precursor ions was set to 30 s.^42^

##### Competition pulldown assay protein identification and quantification

Protein identification and quantification was performed using MaxQuant (v 1.6.1.0)^43^ by searching the LC-MS/MS data against all canonical protein sequences as annotated in the Swissprot reference database (v03.12.15, 20193 entries, downloaded 22.03.2016) using the embedded search engine Andromeda. Carbamidomethylated cysteine was set as fixed modification and oxidation of methionine and N-terminal protein acetylation as variable modifications. Trypsin/P was specified as the proteolytic enzyme and up to two missed cleavage sites were allowed. Precursor tolerance was set to 10 ppm and fragment ion tolerance to 20 ppm. The minimum length of amino acids was set to seven and all data were adjusted to 1% PSM and 1% protein FDR. Label-free quantification^43^ and match between runs was enabled (except for search of experiment corresponding to Fig S1d).^42^

##### Competition pulldown assay data analysis

Relative residual binding of proteins to the affinity matrix was calculated based on the protein intensity ratio relative to the DMSO control for every single inhibitor concentration. EC_50_ values were derived from a four-parameter log-logistic regression using an internal R script that utilizes the ‘drc’ package in R. Targets of the inhibitors were annotated manually. A protein was considered a target or interactor of a target if the resulting binding curve showed a sigmoidal curve shape with a dose dependent decrease of binding to the beads. Additionally, the number of unique peptides and MSMS counts per condition were taken into account.

##### Cell growth assay

Cell confluence (phase) from RPE1 or U2OS cell lines were monitored over time with the IncuCyte S3 (Sartiorius, Germany) in 384-well plates in 50 ul DMEM media, supplemented with 10% Fetal Bovine Serum (FBS) 1% penicillin/Streptomycin and incubated 24h before treatment. Experiment was performed as described in Cano-Franco et al.^44^ 50μl of media containing either 2x final concentration of indicated compounds in or control compounds (final conc.: 0.1% DMSO, 250 nM Torin1, 200ng/mL Bafilomycin) were added and images were taken every two hours over 72 hours. Cell confluence is represented as % of covered area by cells. Each data point represents the averaged ratio or confluence obtained from three individual wells of the plate.

##### Virtual screening

Our compound library of ∼7500 compounds were virtually screened against LC3A and GABARAP (PDB: 6TBE, 4XC2) utilizing SeeSAR (BioSolveIT) and in-house software based upon AutoDock-GPU.^38^ Resulting poses were sorted by estimated affinity (SeeSAR) and free energy (Autodock)^38^ results and filtered by molecular mass with a 750 Da cut-off. 271 compounds were subjected to in-vitro hit validation.

##### Similarity search

Validated hits were included in a Tanimoto-based similarity search via the SpaceLight chemical space exploration tool within the infiniSee suite (BioSolveIT). After removal of already validated hits, 104 additional similars were subjected to *in vitro* analysis.

##### NMR experiments

Prior to measurements, LC3B, GABARAP and GABARAPL2 proteins were equilibrated with buffer containing 25mM HEPES pH=7.0, 100mM NaCl, 5% D2O and 0.15 mM DSS as internal reference. All NMR experiments were performed at a sample temperature of 25 °C on cryogenic probes equipped Bruker Avance spectrometers operating at proton frequencies of 600, 900, and 950 MHz. All NMR spectra were analyzed with the Sparky 3.114 software (University of California, San Francisco, USA). For NMR titration experiments, selected compounds were titrated to 75 μM ^15^N-labeled LC3B, to 50 μM ^13^C,^15^N-labelled GABARAP and to 25 μM ^13^C,^15^N-labelled GABARAPL2 proteins (in standard 5 mm tube, total sample volume 600 μL) to molar ratios of 1:1 and 1:2 (protein:compounds). To achieve reliable calculation of K_D_ values, more titration points were performed for 10O5 and TH152 (up to molar ratio 1:8, proportional to the compounds or complexes solubility). 2D ^1^H-^15^N correlation spectra ([^1^H,^15^N] HSQC for novobiocin, [^15^N,^1^H] BEST-TROSY for other compounds) were recorded at each titration point. CSP values, Δδ, were calculated for each individual backbone amide group using the formula Δδ = [((0.2ΔδN)2 + (ΔδHN)2)/2]1/2 according to the recent guidelines.^45^

##### Crystallization

Initial crystallization hits were obtained by sitting drop vapor diffusion in SwissCi 3-drops plates using a series of commercially available coarse screens. Best hits were obtained in JCSG+ (Hampton Research, USA). Several rounds of optimization were done to meet the conditions required for XChem data collection (high resolution and reproducibility). The test crystals diffracted consistently around 2 Å and as high as 1.36 Å. The selected crystallization condition for further work consisted of 36% PEG 8000 and 0.1 M sodium acetate pH 4.7. For the fragment screening at XChem, the crystals were grown on-site using sitting drop vapor diffusion and the selected condition.

##### Fragment Screening and Structure Solution

A total of 808 fragments from the DSI poised library^46^ (stocks dissolved in DMSO) were transferred to the LC3B crystallization drops using an ECHO liquid handler (20% final DMSO concentration) and soaked for 3 hours before harvesting. Data were collected at the Diamond light source beamline I04-1. A total of 827 datasets were collected (including apo crystals), most of which diffracted to about 2 Å.

Data processing was performed using the automated XChem Explorer pipeline.^47^ Fragment hits were identified using the PanDDA algorithm^48^, followed by visual inspection. Refinement was performed using REFMAC.^49^

