## Supplemental Information for "Targeting LC3/GABARAP for degrader development and autophagy modulation"

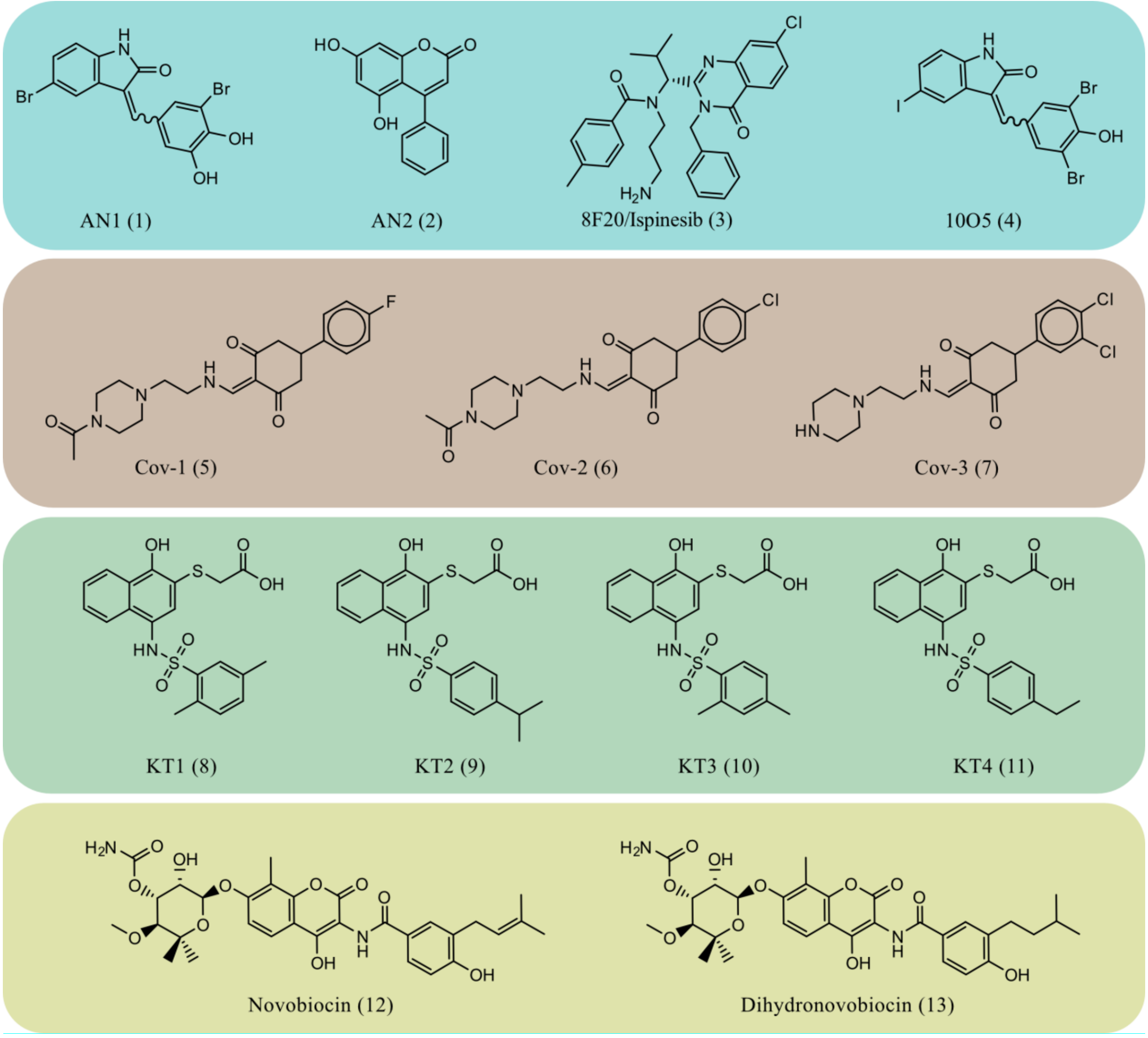


**Supplementary Figure S1:** Chemical structures of the compounds used in this study. Cyan panel: prototypical ATTEC compounds published by Li et al.[1] Brown panel: Derivatives of covalent inhibitors reported by Fan et al.[2] Green panel: Derivatives of compounds reported by Tsuganezawa et al.[3] Yellow panel: Novobiocin and derivative reported by Hartmann et al.[4]

**A**

| ΔTmB [°C] | p62-LIR | NIX-P2-LIR | AN1 (1) | AN2 (2) | 8F20 (3) | 10O5 (4) | Novobiocin (12) |
|---|---|---|---|---|---|---|---|
| LC3A | 1.15 ± 0.04 | 2.39 ± 0.06 | 0.71 ± 0.23 | 0.63 ± 0.09 | -0.43 ± 0.03 | 0.01 ± 0.20 | 0.80 ± 0.30 |
| LC3B | 1.00 ± 0.06 | 1.8 ± 0.17 | 0.69 ± 0.05 | 0.68 ± 0.22 | -0.63 ± 0.06 | 0.43 ± 0.19 | 0.10 ± 0.66 |
| LC3C | 2.09 ± 0.15 | 4.35 ± 0.53 | 0.83 ± 0.21 | 0.33 ± 0.35 | -0.41 ± 0.06 | 0.88 ± 0.29 | 0.03 ± 0.25 |
| GABARAP | 1.31 ± 0.18 | 3.12 ± 0.21 | 0.88 ± 0.10 | 0.04 ± 0.03 | -0.72 ± 0.04 | 0.53 ± 0.16 | 0.13 ± 0.12 |
| GABARAPL1 | 1.18 ± 0.11 | 3.41 ± 0.32 | 0.87 ± 0.21 | 0.83 ± 0.40 | 0.00 ± 0.34 | 1.62 ± 0.45 | 0.53 ± 0.15 |
| GABARAPL2 | 0.92 ± 0.07 | 3.56 ± 0.07 | 0.77 ± 0.53 | 0.32 ± 0.07 | -0.65 ± 0.41 | 1.32 ± 0.91 | 0.30 ± 0.10 |

**B**

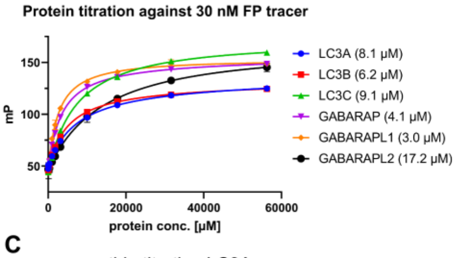


| $K_i$ [µM] | LC3A | LC3B | LC3C | GABARAP | GABARAPL1 | GABARAPL2 |
|---|---|---|---|---|---|---|
| p62-LIR | 15.6 | 10.4 | 26.1 | 6.9 | 5.5 | 32.9 |
| NIX-P2-LIR | 9.7 | 16.1 | 7.5 | 4.9 | 3.7 | 6.3 |
| LIR28 | 503.1 | 852.9 | 197.7 | 1.9 | 1.9 | 5.6 |
| PCM1-LIR | 179.0 | 299.0 | 84.3 | 10.1 | 11.2 | 26.2 |
| AMBRA1-LIR | 1003.8 | 1199.8 | 345.6 | 164.3 | 163.4 | 611.1 |

**C**

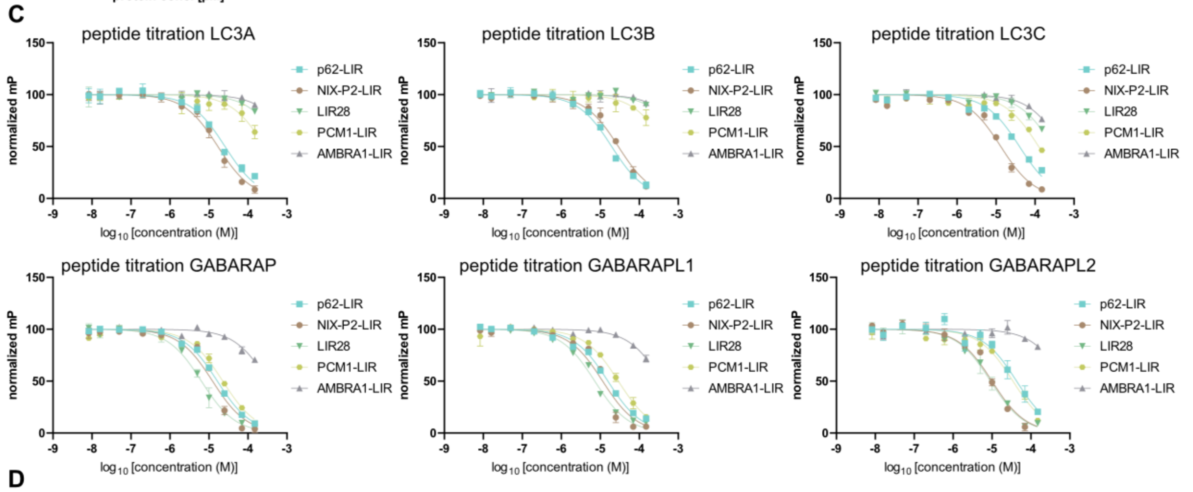


**D**

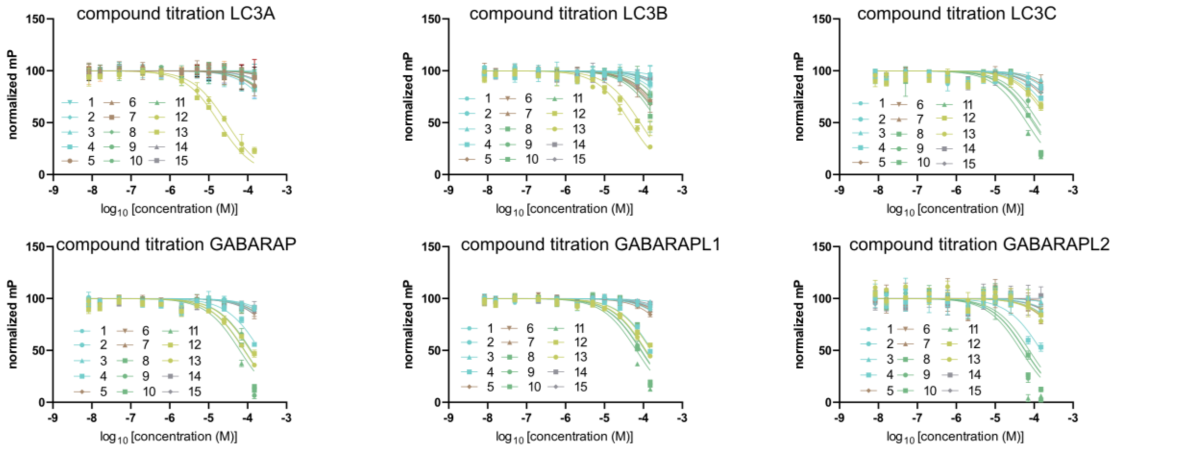


**Supplementary Figure S2:** Establishment of diverse binding assays for all human Atg8 homologs (LC3/GABARAPs). **A)** Results from DSF screening given as °C difference (according to Boltzmann fit) to the DMSO control (ΔTmB). Control peptides p62-LIR and NIX-P2-LIR stabilized the protein up to 4 °C, while compound-induced stabilization could not be observed, also for the positive control compound Novobiocin. Data were measured in technical replicates (n=3). **B)** Protein titration of the different LC3/GABARAPs against the Cy5-labelled p62-LIR peptide depicted with the calculated KD value (left). Table with measured Ki values for a set of different peptides (right), with corresponding curves shown in C). Data were measured in technical replicates (n=3) with error bars depicting the SD. Measured affinities are in agreement with the published affinity data.[5-9] **C)** Peptide displacement experiments of five control peptides against all LC3/GABARAPs. Data were measured in technical replicates (n=2) with error bars depicting the SD. **D)** Displacement curves of the compound set (**1-13**, SI Figure 1) against all LC3/GABARAPs measured in technical replicates (n=2) with error bars depicting the SD.

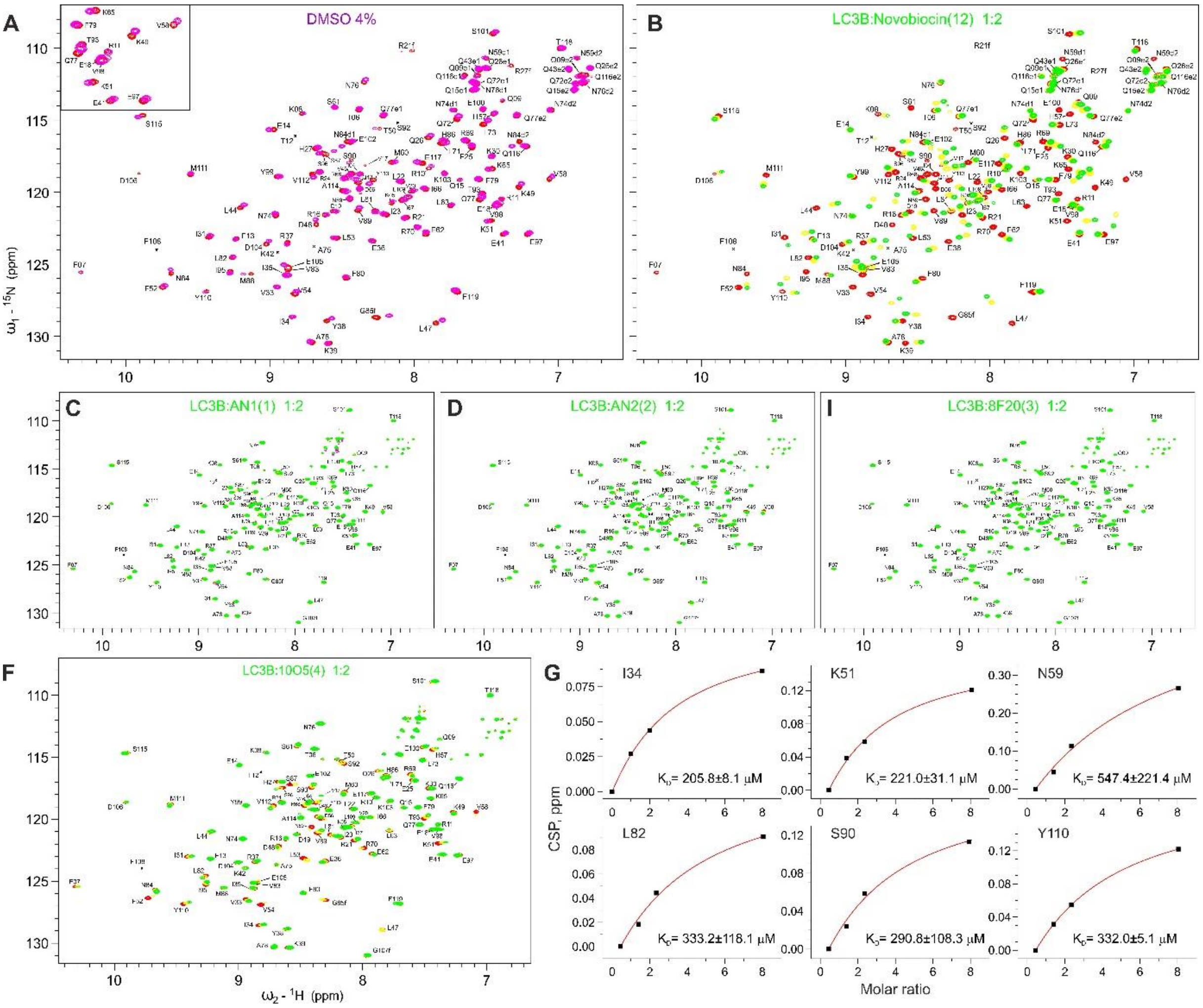


**Supplementary Figure S3:** Interaction between LC3B and compounds **12** and **1**-**4** investigated by NMR. **A)** CSPs of LC3B resonances induced by addition of 4% DMSO into $^{15}$N-labelled LC3B. Reference LC3B [$^{15}$N,$^{1}$H] BEST-TROSY spectrum (red) in overlay with the LC3B spectrum in presence of 4% DMSO (magenta). The small square in the upper left corner shows the LC3B representative (fingerprint) region around residues K51 and V58 depicted in Figure 2D. **C)** NMR titration of LC3B with positive control compound Novobiocin (**12**). Full [$^{1}$H-$^{15}$N]-HSQC spectra of free LC3B (red) overlaid with LC3B spectra in presence of Novobiocin (**12**) in 1:1 molar ratio (yellow) and 1:2 molar ratio (green). **D-F)** NMR titrations of LC3B with AN1 (**1**), AN2 (**2**) and 8F20 (**3**) in 1:1 molar ratio (yellow) and 1:2 molar ratio (green). **G)** NMR titration of LC3B with 10O5 (**4**) in 1:1 molar ratio (yellow) and 1:2 molar ratio (green). **H)** Estimation of $K_D$ values for LC3B:10O5 interaction. $K_D$ values (in µM) calculated for the selected LC3B residues (indicated on each plot). Original CSP values are shown as black squares and the resulting fit is given as a red line in each plot. The selection criteria were I) the residues should be located within or near the interaction surfaces, II) the resonances should be clearly trackable at all titration steps.

**A**

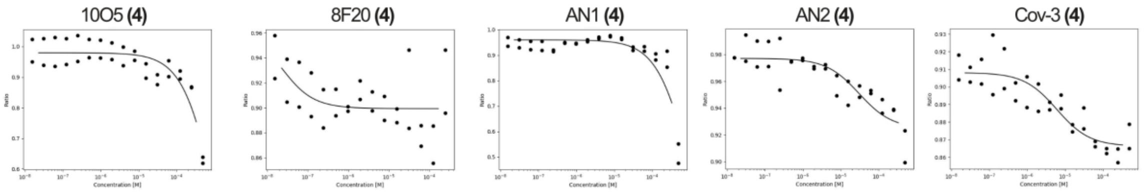


| Ligand | Replicate (n) | Category | Saturation | Δ Ratio | S/N | Kd |
|---|---|---|---|---|---|---|
| 10O5 (**4**) | 2 | Potential Binder | 30.9 % | - 0.979 | 22.9 | - |
| 8F20 (**3**) | 2 | Non-binder | 100.0 % | - 0.064 | 3.1 | - |
| AN1 (**1**) | 2 | Potential Binder | 34.2 % | - 0.960 | 18.9 | - |
| AN2 (**2**) | 2 | Non-binder | 97.5 % | - 0.040 | 4.3 | - |
| Cov-3 (**7**) | 2 | Binder | 84.8% | - 0.512 | 10.4 | 6.08 µM |

**B**

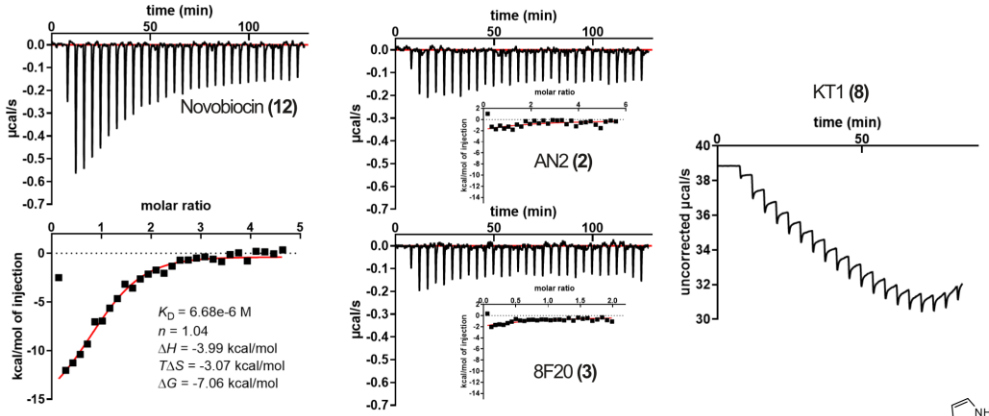


**C**

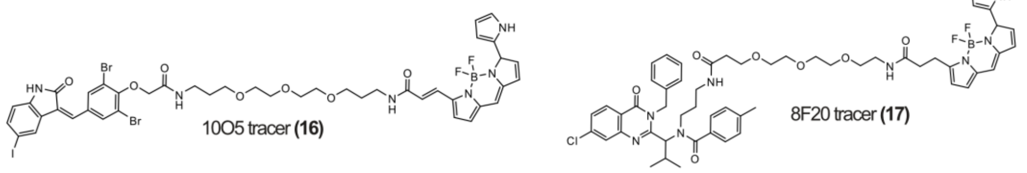


**D**

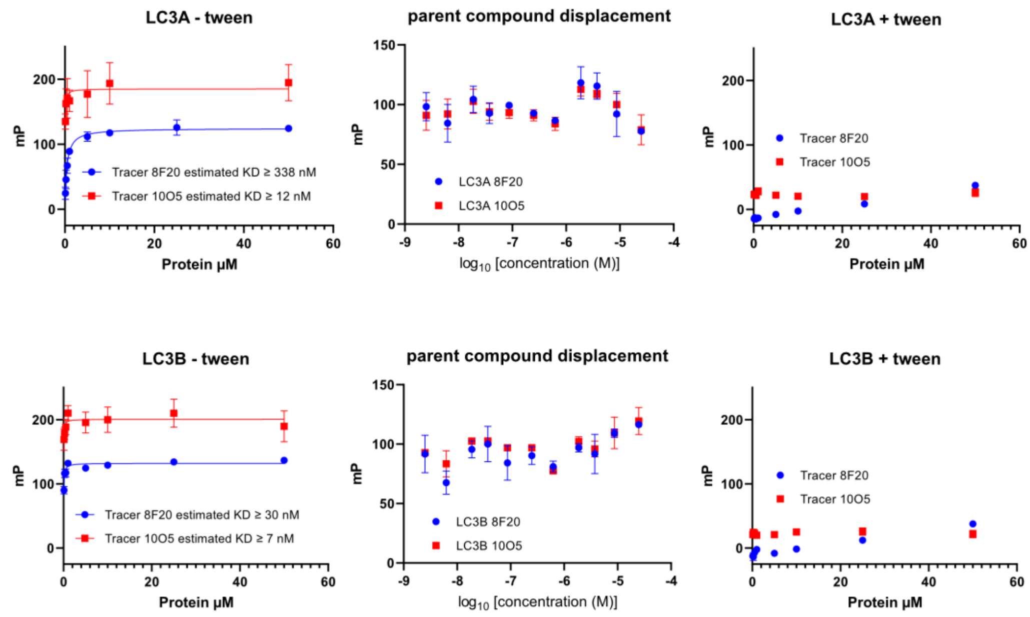


**Supplementary Figure S4: A)** Obtained graphs from the MST measurements of selected compounds (upper panel) and the extracted binding information (lover table) (n=2)**. B)** ITC measurement of Novobiocin (left measurement) and the ATTEC handles AN2 **(2)** and 8F20 **(3)** (middle panels). Residual compounds (**1**, **4**, **5**-**8** etc.) displayed bad solubility in the buffer system and/or led to protein precipitation, resulting in measurements as exemplary depicted for compound **8** (right upper panel). **C)** Structure of the 10O5 (**4**)-based tracer compound **16** and the 8F20 (**3**)-based tracer compound **17**. **D)** Fluorescence polarization assay utilizing the tracers **16** and **17** for interaction studies with LC3A (upper panels) and LC3B (lower panels). The left panels depict protein titration measurements for tracer $K_D$ determination indicating nanomolar tracer affinity as presented in[10], lacking detergent in the buffer to suppress unspecific binding. Middle panels depict displacement experiments using compound **3** for tracer **17** and compound **4** for tracer **16** showing no displacement, indicating unspecific binding during protein titration. Right panels show a repeated protein titration with 0.05% Tween20 supplemented buffer, completely abolishing the putative binding, revealing unspecific binding of the tracers. Data were collected in technical triplicates with error bars expressing the SD (n=3).

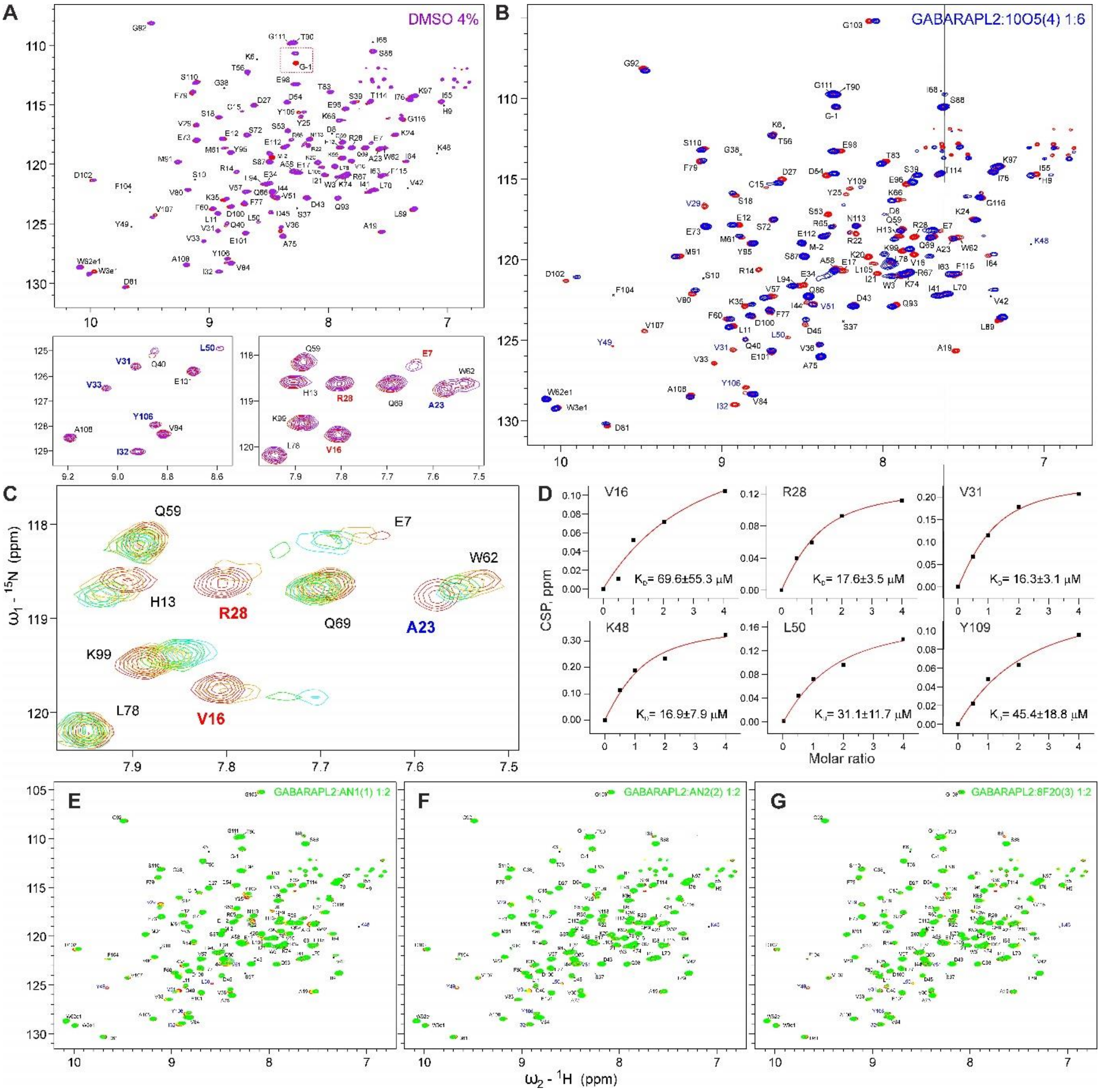


**Supplementary Figure S5:** Interaction between GABARAPL2 and compounds **1**-**4** investigated by NMR. **A)** CSPs of GABARAPL2 resonances induced by addition of 4% DMSO into $^{13}$C,$^{15}$N-labelled GABARAPL2. Upper plot: reference GABARAPL2 [$^{15}$N,$^{1}$H] BEST-TROSY full spectrum (red) in overlay with the GABARAPL2 spectrum in presence of 4% DMSO. Lower plots show the GABARAPL2 representative (fingerprint) regions around residues L50 (left) and V16 (right). **C-E)** NMR titration of GABARAPL2 with 10O5 (**4**). **B)** Full [$^{15}$N,$^{1}$H] BEST-TROSY spectrum of free GABARAPL2 (red) overlaid with GABARAPL2 spectrum in presence of 10O5 in 1:6 molar ratio (blue). **C)** Representative region of the [$^{15}$N,$^{1}$H] BEST-TROSY GABARAPL2 spectrum upon titration of GABARAPL2 with 10O5 (free protein – red, molar ratios of 1:0.5, 1:1, 1:2 and 1:4 are given in orange, yellow, green and cyan, respectively). **D)** Estimation of $K_D$ values for GABARAPL2:10O5 (**4**) interaction. $K_D$ values (in µM) calculated for the selected GABARAPL2 residues (indicated on each plot). Original CSP values are shown as black squares and the resulting fit is given as a red line in each plot. The selection criteria were identical as described in SI Figure S3. **E-G)** NMR titrations of GABARAPL2 with AN1 (**1**), AN2 (**2**) and 8F20 (**3**), respectively.

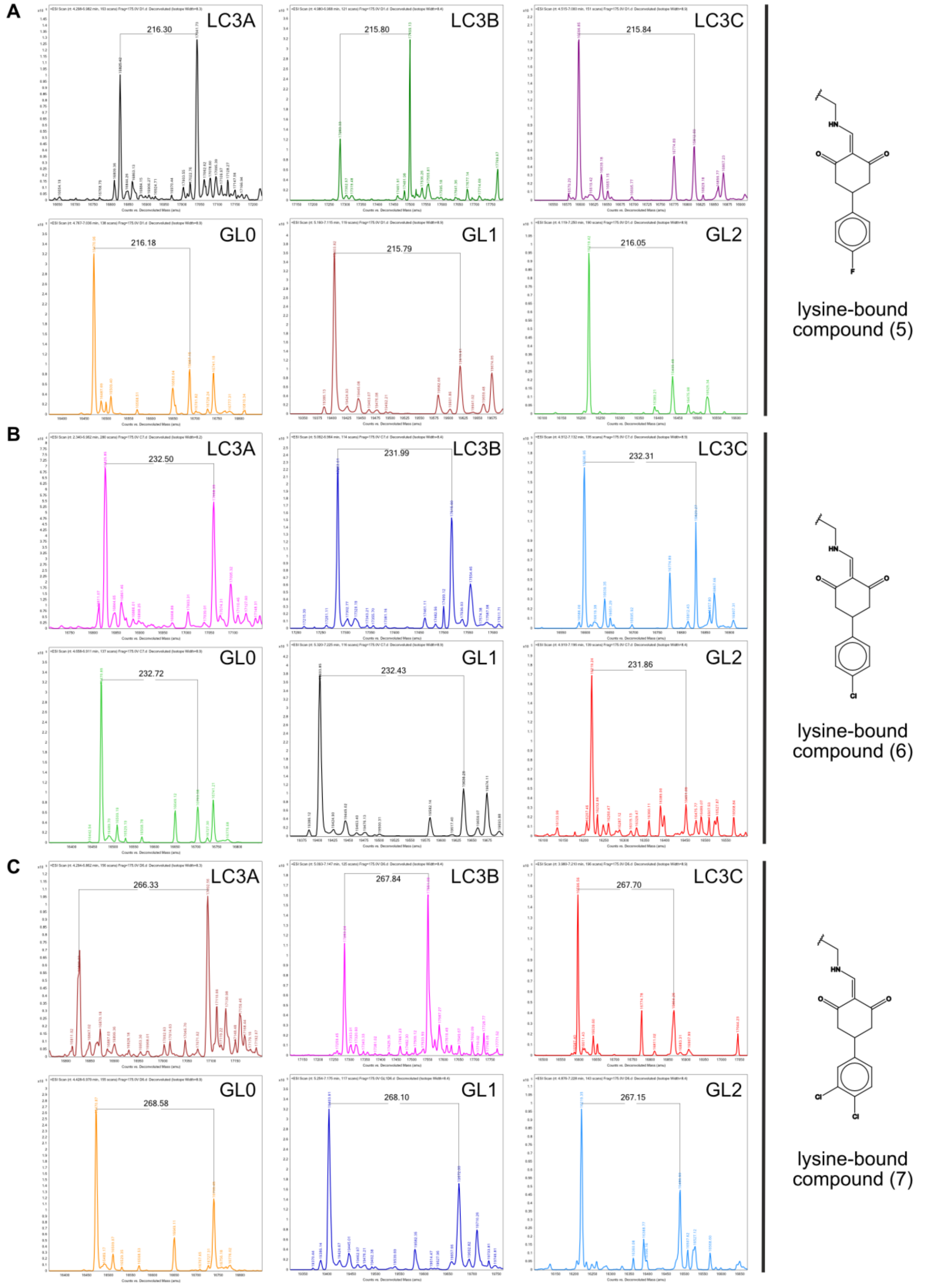


**Supplementary Figure S6:** ESI mass spectrometry for all LC3/GABARAP proteins treated with compounds **5**-**7**. A) Mass shifts obtained from treatment with compound **5**. All LC3/GABARAPs were successfully labelled with the highest labelling ratio on LC3A und LC3B. B) Mass shifts obtained from treatment with compound **6**. All LC3/GABARAPs were successfully labelled with the highest labelling ratio on all LC3 proteins. C) Mass shifts obtained from treatment with compound **7**. All Atg8 homologs were successfully modified with the highest ratio on LC3A und LC3B.

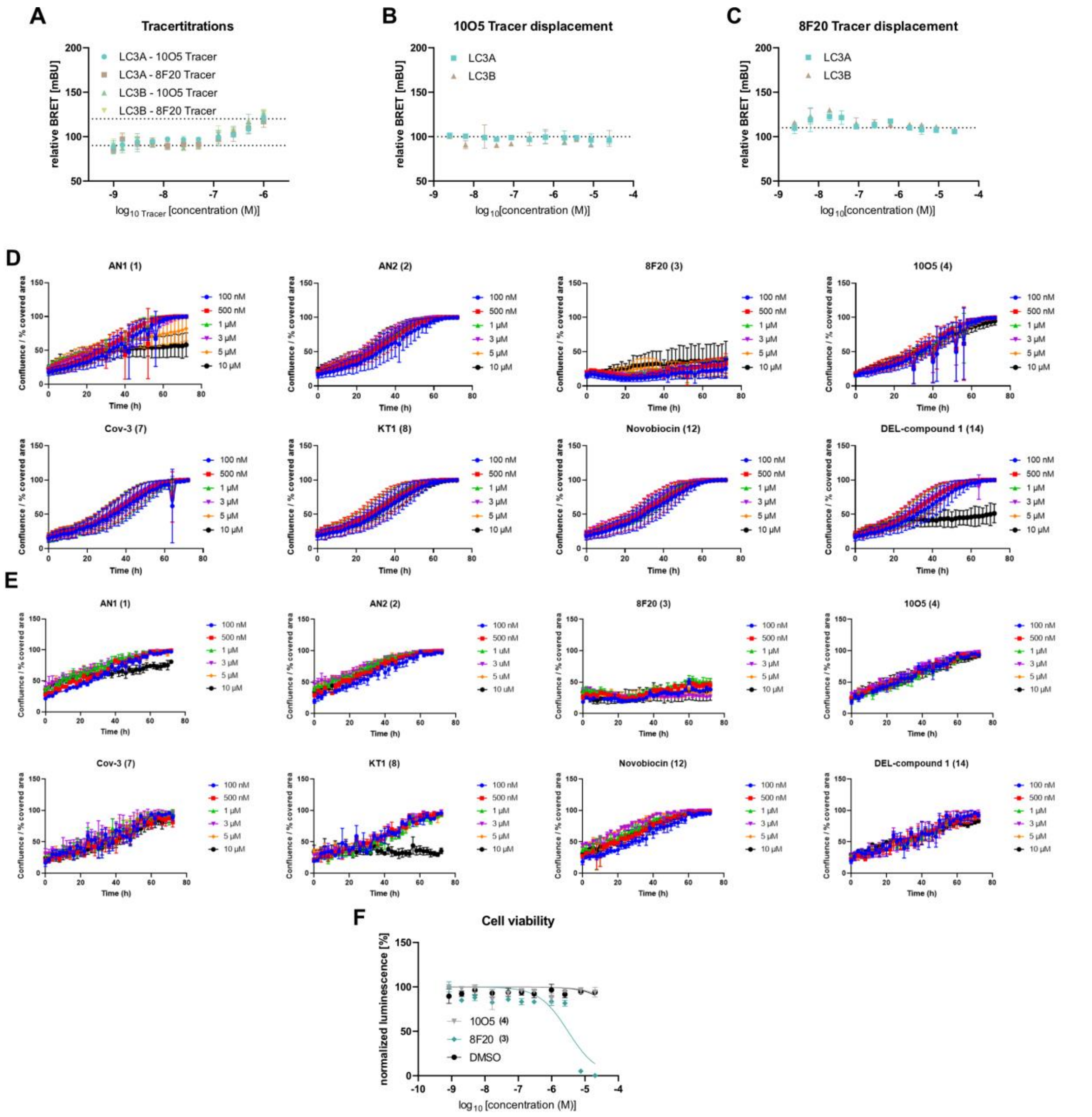


**Supplementary Figure S7: A)** tracer titrations of compounds **16** and **17** (SI Figure S4) against NanoLuciferase-tagged LC3A and LC3B showing a maximum BRET increase of ~1.3-fold, indicating unspecific BRET increase due to the presence of higher fluorophore concentrations. Displacement assays were carried out to test specific compound binding on LC3A (**B)**) and LC3B (**C)**) through tracer displacement using compound **4** for tracer compound **16** and compound **3** for tracer compound **17**. No displacement at 1 µM of the tracer compounds proves the hypothesis of unspecific BRET increase, therefore showing no compound-LC3 interaction. Measurements were carried out in biological replicates (n=2) with error bars expressing the SD. **D)** RPE1 growth was detected based on the cells confluence within the well. Outliers were determined to be faulty collected images. Apart from high dosage effects of compounds **1**, only compound **3** lead to significant growth reduction. Data were collected in biological replicates with technical triplicates each (n=6) and are presented as mean with error bars expressing the SD. **E)** Growth curves of U2OS cells, treated with different small molecules. Growth was detected based on the cells confluence within the well. For U2OS cells, compound **8** shows a drastic effect at the highest concentration, while compound **3** leads to significant growth inhibition in all tested concentrations. Data were collected in technical triplicates (n=3) and error bars expressing the SD. **F)** Cell viability measurements of the ATTEC handles 8F20 (**3**) and 10O5 (**4**) indicating no viability/toxicity effect of compound **4** while compound **3** treatment led to viability loss in the high µM range. Data were collected in biological replicates (n=2) with error bars expressing the SD. Data is in agreement with published viability data.[1]

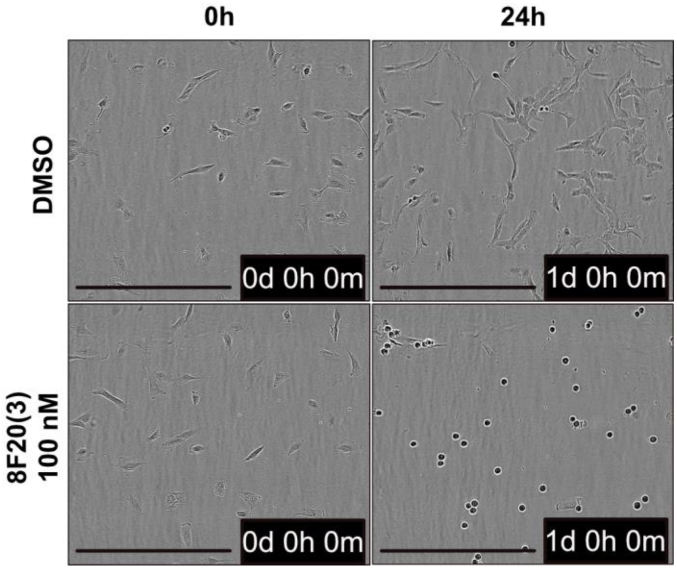


**Supplementary Figure S8:** Microscopy images **acquired with the IncuCyte S3 (10x) show rounding of RPE1 cells** after treatment with **compound** 3 for 24h. Scale bar reflects 500 um.

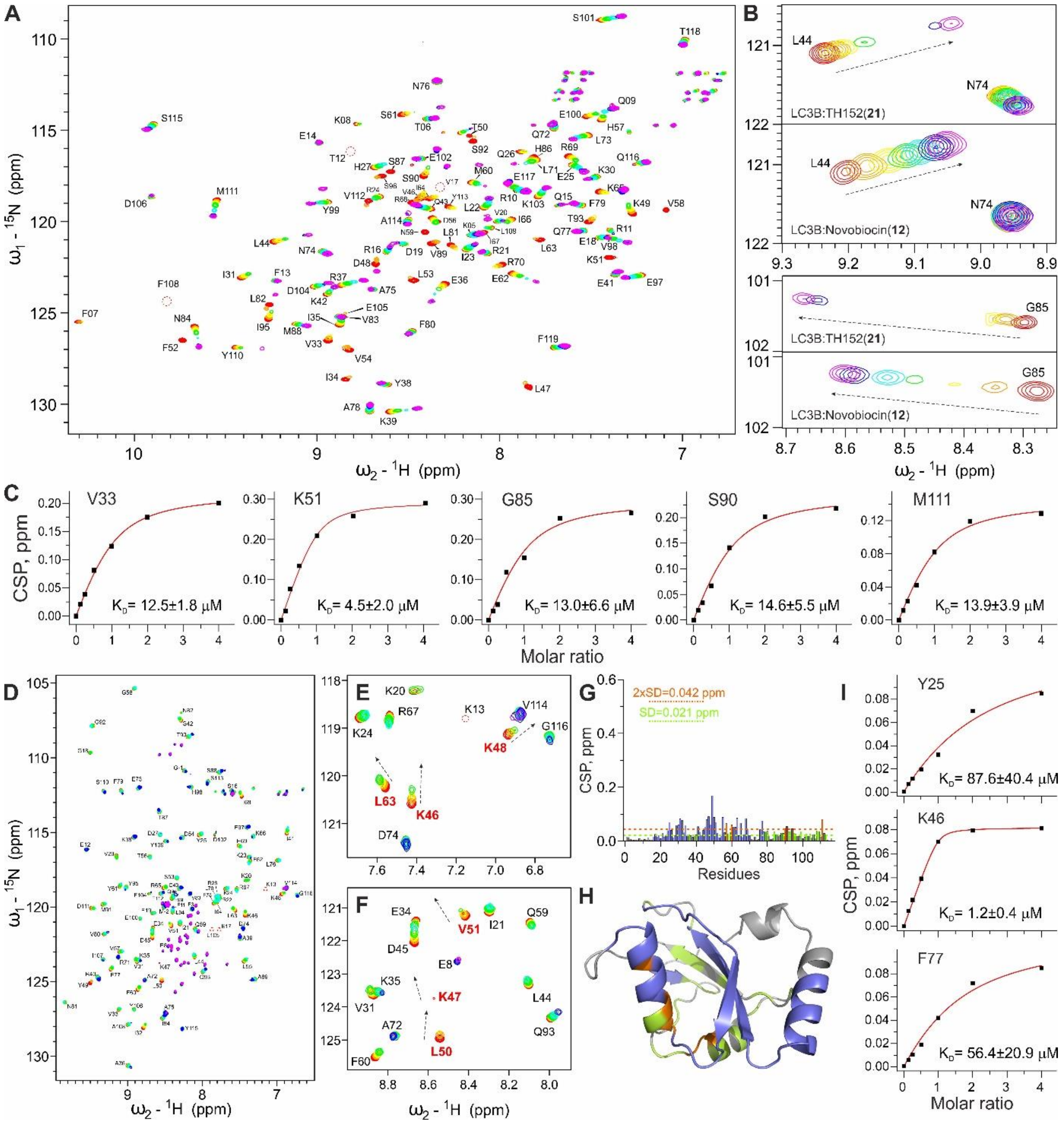


**Supplementary Figure S9: A-D)** Interaction between LC3B and compound TH152 (**21**) investigated by NMR. **A)** Full [$^{15}$N,$^{1}$H] BEST-TROSY spectrum of free LC3B (red) overlaid with LC3B spectra in presence of **21** in increasing molar ratios (1:0.125 – orange, 1:0.25 – yellow, 1:0:5 – green, 1:1 – cyan, 1:2 – blue and 1:4 - magenta). **B)** LC3B:**21** interaction revealed strong similarity with LC3B:**12** interaction. Upper plot: representative spectra area around L44 backbone HN resonance for titration with **21** and **12** (indicated on each graph). Lower plot: representative spectra area around G85 backbone HN resonance for titration with **21** and **12**. **C)** Estimation of $K_D$ values for LC3B:**21** interaction. $K_D$ values calculated for the selected LC3B residues (indicated on each plot) upon titration with **21**. Original CSP values are shown as black squares and the resulting fit is given as a red line in each plot. The selection criteria were identical as described in SI Figure S3. **E-J)** Interaction between GABARAP and compound **21** investigated by NMR. **D)** Full [$^{15}$N,$^{1}$H] BEST-TROSY spectrum of free GABARAP overlaid with GABARAP spectra in presence of **21** in increasing molar ratios (the same colorcode as for LC3B titration). Non-assigned peaks in the presented spectra are from Gln and Asn sidechain $NH_2$ resonances, and from traces of the GABARAP degradation peptides. **E-F)** Representative regions of the [$^{15}$N,$^{1}$H] BEST-TROSY GABARAP spectra around key residues K46, K48 and L63 (**E**), and K47, L50 and V51

(**F**) upon titration of GABARAP with **21** (the same colorcode as above). **G)** CSP values, induced by compounds **21** at molar ratio 1:2, are plotted against GABARAP residue numbers. The light green dashed line indicates the standard deviations (SD) over all residues, the orange dashed line indicates double SD values. Residues with small (CSP < SD), intermediate (SD < CSP < 2xSD) or strong (2xSD < CSP) CSP values are marked in grey, light green and orange, respectively. GABARAP residues which undergo strong intermediate exchange mode (significant decrease of the resonances intensity upon titration with **21**) are marked blue. **H)** 3D mapping of CSP values on GABARAP structure (pdb: 1UGM), indicating both hydrophobic pockets, HP1 and HP2, as most relevant interaction sites. The colorcode is the same as above. **I)** Estimation of $K_D$ values for GABARAP:**21** interaction. $K_D$ values (in µM) calculated for the selected GABARAP residues (indicated on each plot) upon titration with **21**. Original CSP values are shown as black squares and the resulting fit is given as a red line in each plot. The selection criteria were identical as described in SI Figure S3.

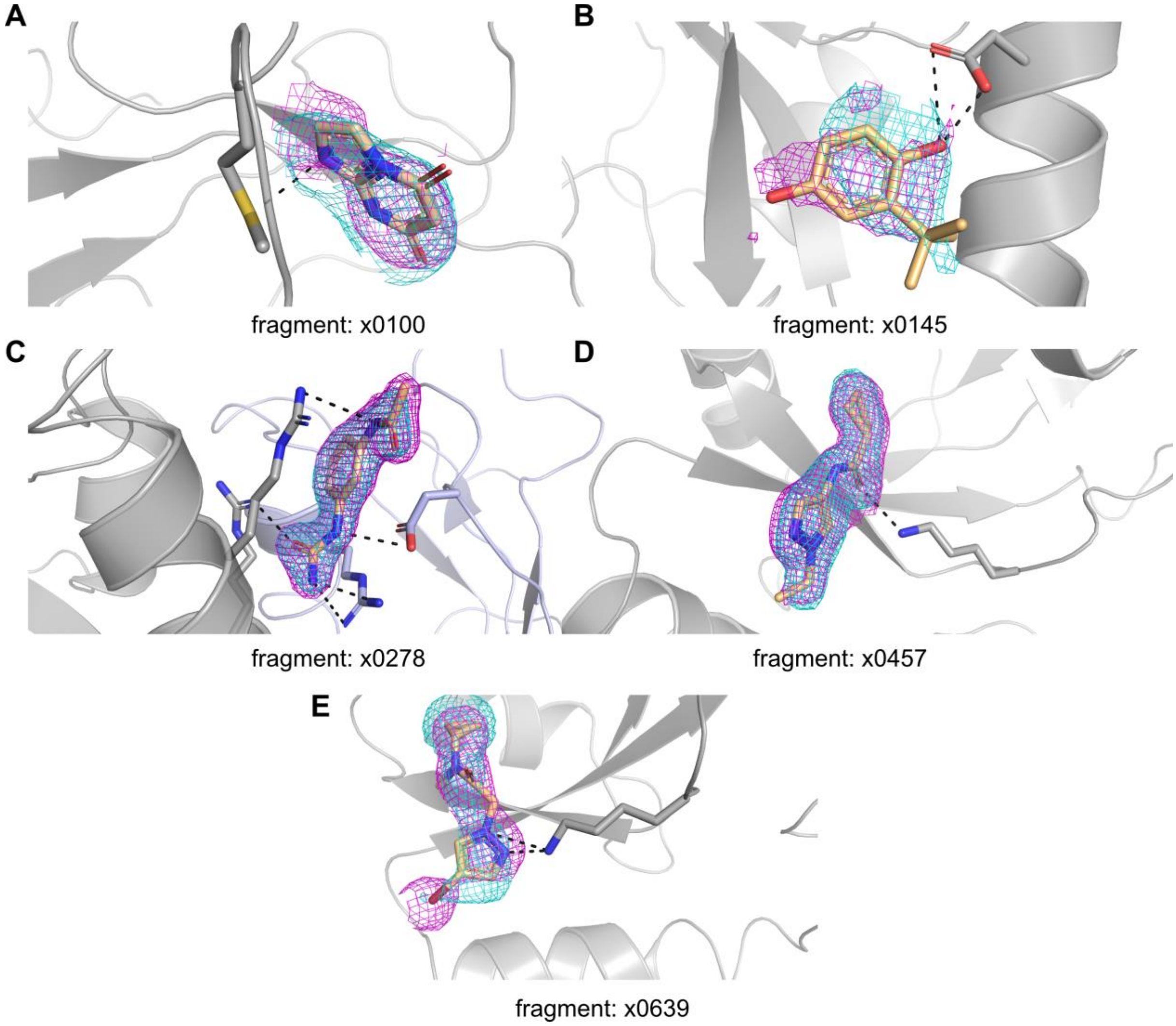


**Supplementary Figure S10:** Crystallized fragments bound to LC3A as depicted in Figure 5. Together with the fragments bound in the crystal structure, the event map (cyan) and the 2Fo-Fc (magenta) is depicted together with interacting residues.

**Table 1:** Structure-activity relationship of sulfonamide based compounds[12-14] on LC3A tracer displacement, normalized on p62-LIR positive control.

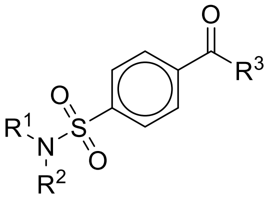

| Cpd-ID | $R^1$ | $R^2$ | $R^3$ | % tracer bound @LC3A |
|---|---|---|---|---|
| (TH061) | 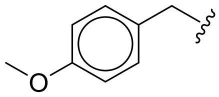 | 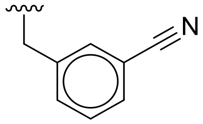 | OH | 83 |
| (TH119) | 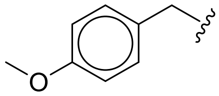 | 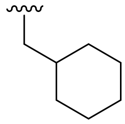 | OH | 92 |
| (TH133) | 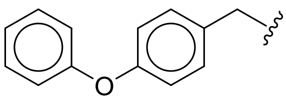 | 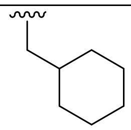 | OH | 82 |
| (TH253) | 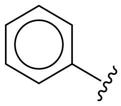 | H | 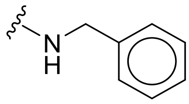 | 88 |
| (TH676) | 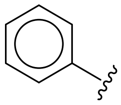 | H | 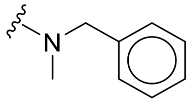 | 84 |
| (TH281) | 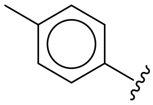 | H | 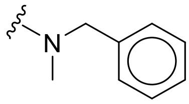 | 95 |
| (TH677) | 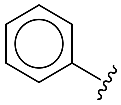 | H | 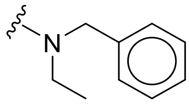 | 87 |

**Table 2:** Structure-activity relationship of benzothiazole based compounds[12,15] on LC3A tracer displacement, normalized on p62-LIR positive control.

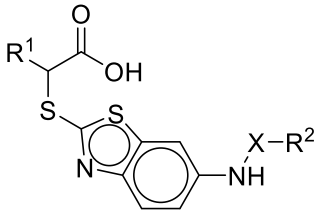


| Cpd-ID | $R^1$ | $R^2$ | X | % tracer bound @LC3A |
|---|---|---|---|---|
| (TH195) | *n*-hexyl | 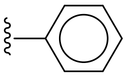 | $CH_2$ | 78 |
| (TH185) | *n*-hexyl | 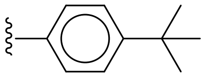 | $CH_2$ | 82 |
| (TH198) | *n*-hexyl | 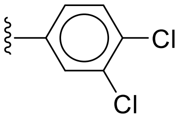  | $CH_2$ | 79 |
| (TH191) | *n*-hexyl | 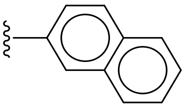 | $CH_2$ | 78 |
| (TH200) | *n*-hexyl | 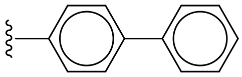 | $CH_2$ | 72 |
| (TH181) | *n*-hexyl | 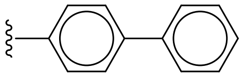 | C=O | 78 |
| (TH177) | *n*-hexyl | 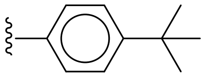 | C=O | 85 |
| (TH188) | *n*-butyl | 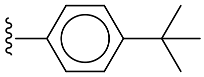 | C=O | 89 |

**Table 3:** Structure-activity relationship of pyrimidine based compounds[12,16] on LC3A tracer displacement, normalized on p62-LIR positive control.

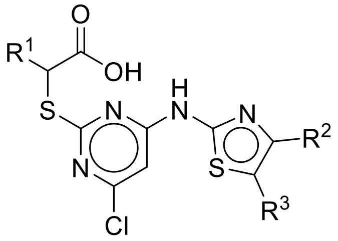


| Cpd-ID | $R^1$ | $R^2$ | $R^3$ | % tracer bound @LC3A |
|---|---|---|---|---|
| (TH020) | *n*-hexyl | 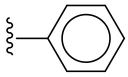 | H | 84 |
| (TH018) | *n*-hexyl | 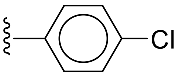 | H | 81 |
| (TH035) | ethyl | 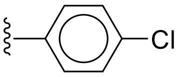 | H | 85 |
| (TH026) | *n*-hexyl | 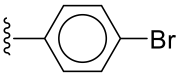 | H | 79 |
| (TH057) | *n*-hexyl | 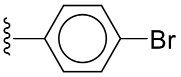 | Methyl | 94 |
| (TH100) | *n*-hexyl | 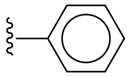 | Ethyl | 77 |
| (TH083) | *n*-hexyl | 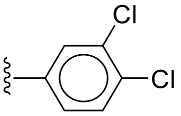 | H | 84 |
| (TH102) | *n*-hexyl | 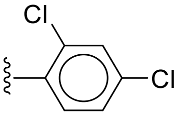 | H | 78 |
| (TH044) | *n*-hexyl | 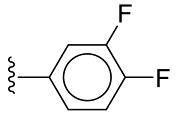 | H | 76 |
| (TH054) | *n*-hexyl | 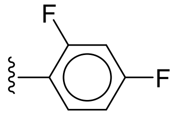 | H | 75 |
| (TH084) | *n*-hexyl | 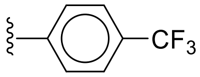 | H | 76 |
| (TH107) | ethyl | 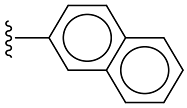 | H | 82 |
| (TH034) | *n*-hexyl | 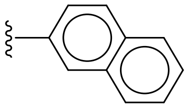 | H | 79 |
| (TH104) | *n*-hexyl | 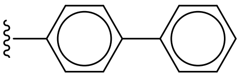 | H | 82 |
| (TH105) | *n*-hexyl | 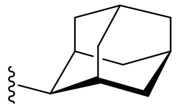 | H | 84 |

| | | | | |
|---|---|---|---|---|
| (TH062) | *n*-hexyl | 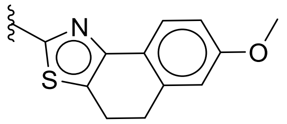 | | 76 |
| (TH121) | *n*-hexyl | 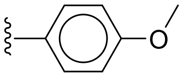 | H | 83 |
| (TH056) | *n*-hexyl | 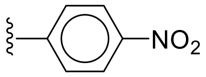 | H | 61 |
| **21 (TH152)** | *n*-hexyl | 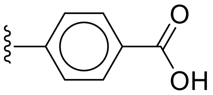 | H | 26 |

Table 4: X-ray data collection and refinement statistics

| | 8Q53 | 7GAU | 7GA8 | 7GA9 | 7GAA | 7GAB |
|---|---|---|---|---|---|---|
| **Data collection**[a] | | | | | | |
| Wavelength (Å) | 0.99999 | 0.92124 | 0.92124 | 0.92124 | 0.92124 | 0.92124 |
| Space group | P43 | P43 | P43 | P43 | P43 | P43 |
| Resolution (Å) | 42.61 – 1.36 | 61.00 – 1.59 | 30.68 – 1.87 | 30.49 – 2.17 | 30.91 – 2.03 | 43.05 – 2.23 |
| Last resolution shell (Å) | 1.38 – 1.36 | 1.62 – 1.59 | 1.91 – 1.87 | 2.23 – 2.17 | 2.08 – 2.03 | 2.31 – 2.23 |
| Unit cell parameters | | | | | | |
| a,b,c (Å) | 60.26, 60.26, 34.93 | 61.01, 61.01, 35.48 | 60.83, 60.83, 35.53 | 60.98, 60.98, 35.32 | 60.59, 60.59, 35.93 | 60.89, 60.89, 35.59 |
| α, β, γ (°) | 90, 90, 90 | 90, 90, 90 | 90, 90, 90 | 90, 90, 90 | 90, 90, 90 | 90, 90, 90 |
| Total number of observations | 181809 (17402) | 241587 (10881) | 150104 (9935) | 93140 (6872) | 116879 (8961) | 87203 (7854) |
| Unique reflections | 26382 (2575) | 17826 (896) | 10871 (687) | 7021 (515) | 8621 (641) | 6512 (584) |
| Mosaicity (˚) | 0.13 | 0 | 0 | 0 | 0 | 0 |
| Multiplicity | 6.9 (6.8) | 13.6 (12.1) | 13.8 (14.5) | 13.3 (13.3) | 13.6 (14.0) | 13.4 (13.4) |
| Mean I/σ(I) | 21.11 (2.40) | 11.9 (0.3) | 14.4 (0.2) | 8.8 (0.9) | 12.3 (0.4) | 10.3 (0.5) |
| Completeness (%) | 97.08 (96.12) | 100.0 (100.0) | 99.5 (100.0) | 99.7 (98.4) | 100.0 (100.0) | 100.0 (100.0) |
| $R_{merge}$[b] | 0.042 (0.733) | 0.092 (3.870) | 0.083 (4.560) | 0.199 (3.303) | 0.186 (10.910) | 0.144 (3.603) |
| $R_{meas}$[c] | 0.046 (0.794) | 0.095 (1.040) | 0.087 (4.725) | 0.208 (3.433) | 0.194 (11.319) | 0.150 (3.747) |
| $R_{pim}$[d] | 0.017 (0.302) | 0.026 (1.150) | 0.024 (1.235) | 0.058 (0.931) | 0.053 (3.000) | 0.041 (1.022) |
| **Refinement** | | | | | | |
| Resolution (Å) | 42.61 – 1.36 | 61.00 – 1.59 | 30.68 – 1.87 | 30.49 – 2.17 | 30.91-2.03 | 43.05 – 2.23 |
| Reflections used | 26378 | 16693 | 10093 | 6749 | 8268 | 6243 |
| Free R flagged reflections | 1934 | 780 | 426 | 266 | 330 | 251 |
| $R_{cryst}$[e] | 0.190 | 0.193 | 0.216 | 0.180 | 0.190 | 0.187 |
| $R_{free}$[f] | 0.210 | 0.215 | 0.274 | 0.246 | 0.248 | 0.276 |
| rmsd bonds (Å) | 0.007 | 0.008 | 0.006 | 0.008 | 0.007 | 0.009 |
| rmsd angles (˚) | 0.99 | 1.542 | 1.428 | 1.518 | 1.547 | 1.625 |
| Ramachandran plot | | | | | | |
| Most favored (%) | 99.12 | 99.12 | 99.12 | 99.12 | 99.12 | 97.35 |
| Additionally allowed (%) | 0.88 | 0.88 | 0.88 | 0.88 | 0.88 | 2.65 |
| Mean B-factor (Å$^2$) | 25.830 | 36.989 | 55.917 | 56.978 | 65.188 | 66.317 |

[a]Values for the last resolution shell are in parentheses.

[b]$R_{merge} = \Sigma_{hkl}\Sigma_i|I_i(hkl)-\langle I(hkl)\rangle| / \Sigma_{hkl}\Sigma_i I_i(hkl)$, where I(hkl) is the intensity of reflection hkl

[c]$R_{meas} = \Sigma_{hkl}(n/(n-1))^{1/2}\Sigma_i|I_i(hkl)-\langle I(hkl)\rangle| / \Sigma_{hkl}\Sigma_i I_i(hkl)$

[d]$R_{pim} = \Sigma_{hkl}(1/(n-1))^{1/2}\Sigma_i|I_i(hkl)-\langle I(hkl)\rangle| / \Sigma_{hkl}\Sigma_i I_i(hkl)$

[e]$R_{cryst} = \Sigma_{hkl} ||F_{obs}|-|F_{calc}|| / \Sigma|F_{obs}|$

[f]$R_{free}$ is the cross-validation R-factor computed for the test set of unique reflections.

| | 7GAC | 7GAD | 7GAE | 7GAF | 7GAG | 7GAH |
|---|---|---|---|---|---|---|
| **Data collection**[a] | | | | | | |
| Wavelength (Å) | 0.92124 | 0.92124 | 0.92124 | 0.92124 | 0.92124 | 0.92124 |
| Space group | P43 | P43 | P43 | P43 | P43 | P43 |
| Resolution (Å) | 42.88 – 1.91 | 61.02 – 1.86 | 43.08 – 1.92 | 42.98 – 1.84 | 61.00 – 1.59 | 30.60 – 1.90 |
| Last resolution shell (Å) | 1.95 – 1.91 | 1.90 – 1.86 | 1.96 – 1.92 | 1.88 – 1.84 | 1.62 – 1.59 | 1.95 – 1.90 |
| Unit cell parameters | | | | | | |
| a,b,c (Å) | 60.62, 60.62, 35.85 | 61.03, 61.03, 35.55 | 60.92, 60.92, 35.65 | 60.80, 60.80, 35.38 | 61.01, 61.01, 35.48 | 61.20, 61.20, 35.27 |
| α, β, γ (°) | 90, 90, 90 | 90, 90, 90 | 90, 90, 90 | 90, 90, 90 | 90, 90, 90 | 90, 90, 90 |
| Total number of observations | 140194 (9706) | 152869 (9459) | 139496 (9520) | 156378 (9812) | 241587 (10881) | 138579 (11039) |
| Unique reflections | 10254 (684) | 11186 (665) | 10230 (675) | 11438 (689) | 17826 (896) | 10488 (771) |
| Mosaicity (˚) | 0 | 0 | 0 | 0 | 0 | 0 |
| Multiplicity | 13.7 (14.2) | 13.7 (14.2) | 13.6 (14.1) | 13.7 (14.2) | 13.6 (12.1) | 13.2 (14.3) |
| Mean I/σ(I) | 9.8 (0.3) | 15.2 (0.2) | 11.6 (0.4) | 13.4 (0.3) | 11.9 (0.3) | 15.0 (1.0) |
| Completeness (%) | 99.5 (100.0) | 99.8 (96.4) | 99.8 (97.5) | 100.0 (99.7) | 100.0 (100.0) | 100.0 (100.0) |
| $R_{merge}$[b] | 0.125 (5.485) | 0.082 (5.795) | 0.113 (3.518) | 0.088 (5.554) | 0.092 (3.870) | 0.127 (2.878) |
| $R_{meas}$[c] | 0.131 (5.665) | 0.086 (6.163) | 0.118 (3.652) | 0.091 (5.761) | 0.095 (1.040) | 0.133 (2.985) |
| $R_{pim}$[d] | 0.036 (1.506) | 0.023 (1.585) | 0.045 (1.378) | 0.025 (1.526) | 0.026 (1.150) | 0.038 (0.787) |
| **Refinement** | | | | | | |
| Resolution (Å) | 42.88 – 1.91 | 61.02 – 1.86 | 43.08 – 1.92 | 42.98 – 1.84 | 61.00 – 1.59 | 30.60 – 1.90 |
| Reflections used | 9760 | 10362 | 9804 | 10740 | 16693 | 10053 |
| Free R flagged reflections | 408 | 436 | 407 | 468 | 780 | 421 |
| $R_{cryst}$[e] | 0.207 | 0.201 | 0.197 | 0.194 | 0.193 | 0.199 |
| $R_{free}$[f] | 0.260 | 0.258 | 0.251 | 0.245 | 0.215 | 0.254 |
| rmsd bonds (Å) | 0.008 | 0.008 | 0.008 | 0.009 | 0.008 | 0.009 |
| rmsd angles (˚) | 1.539 | 1.524 | 1.594 | 1.591 | 1.542 | 1.628 |
| Ramachandran plot | | | | | | |
| Most favored (%) | 100 | 99.12 | 99.12 | 100 | 99.12 | 100 |
| Additionally allowed (%) | 0.00 | 0.88 | 0.88 | 0.00 | 0.88 | 0.00 |
| Mean B-factor (Å$^2$) | 53.402 | 53.986 | 50.216 | 53.317 | 36.989 | 47.276 |

[a]Values for the last resolution shell are in parentheses.

[b]$R_{merge} = \Sigma_{hkl}\Sigma_i|I_i(hkl)-\langle I(hkl)\rangle| / \Sigma_{hkl}\Sigma_i I_i(hkl)$, where I(hkl) is the intensity of reflection hkl

[c]$R_{meas} = \Sigma_{hkl}(n/(n-1))^{1/2}\Sigma_i|I_i(hkl)-\langle I(hkl)\rangle| / \Sigma_{hkl}\Sigma_i I_i(hkl)$

[d]$R_{pim} = \Sigma_{hkl}(1/(n-1))^{1/2}\Sigma_i|I_i(hkl)-\langle I(hkl)\rangle| / \Sigma_{hkl}\Sigma_i I_i(hkl)$

[e]$R_{cryst} = \Sigma_{hkl} ||F_{obs}|-|F_{calc}|| / \Sigma|_{Fobs}|$

[f]$R_{free}$ is the cross-validation R-factor computed for the test set of unique reflections.

| | 7GAI | 7GAJ | 7GAK | 7GAL | 7GAM | 7GAN |
|---|---|---|---|---|---|---|
| **Data collection**[a] | | | | | | |
| Wavelength (Å) | 0.92124 | 0.92124 | 0.92124 | 0.92124 | 0.92124 | 0.92124 |
| Space group | P43 | P43 | P43 | P43 | P43 | P43 |
| Resolution (Å) | 61.00 – 1.97 | 43.15 – 1.89 | 30.40 – 1.77 | 43.05 – 1.90 | 60.67 – 1.75 | 43.19 – 2.09 |
| Last resolution shell (Å) | 2.02 – 1.97 | 1.93 – 1.89 | 1.81 – 1.77 | 1.94 – 1.90 | 1.78 – 1.75 | 2.15 – 2.09 |
| Unit cell parameters | | | | | | |
| a,b,c (Å) | 60.98, 60.98, 35.59 | 61.00, 61.00, 35.33 | 60.81, 60.81, 35.10 | 60.87, 60.87, 35.41 | 60.68, 60.68, 35.61 | 61.09, 61.09, 35.54 |
| α, β, γ (°) | 90, 90, 90 | 90, 90, 90 | 90, 90, 90 | 90, 90, 90 | 90, 90, 90 | 90, 90, 90 |
| Total number of observations | 128225 (9138) | 145177 (9643) | 173163 (9502) | 141587 (9231) | 180563 (9230) | 106979 (8388) |
| Unique reflections | 9444 (650) | 10634 (680) | 12673 (736) | 10401 (656) | 13277 (723) | 7949 (608) |
| Mosaicity (˚) | 0 | 0 | 0 | 0 | 0 | 0 |
| Multiplicity | 13.6 (14.1) | 13.7 (14.2) | 13.7 (12.9) | 13.6 (14.1) | 13.6 (12.8) | 13.5 (13.8) |
| Mean I/σ(I) | 14.7 (0.4) | 10.5 (0.3) | 13.6 (0.3) | 9.2 (0.3) | 14.3 (0.4) | 10.4 (0.5) |
| Completeness (%) | 99.9 (98.4) | 100.0 (100.0) | 100.0 (99.4) | 99.9 (99.2) | 99.9 (98.3) | 99.9 (99.3) |
| $R_{merge}$[b] | 0.076 (2.872) | 0.113 (5.099) | 0.104 (6.109) | 0.118 (3.840) | 0.068 (2.668) | 0.124 (3.388) |
| $R_{meas}$[c] | 0.079 (2.980) | 0.118 (5.288) | 0.109 (6.361) | 0.123 (3.985) | 0.070 (2.878) | 0.129 (3.517) |
| $R_{pim}$[d] | 0.022 (0.793) | 0.032 (1.395) | 0.030 (1.764) | 0.033 (1.059) | 0.026 (1.122) | 0.035 (0.937) |
| **Refinement** | | | | | | |
| Resolution (Å) | 61.00 – 1.97 | 43.15 – 1.89 | 30.40 – 1.77 | 43.05 – 1.90 | 60.67 – 1.75 | 43.19 – 2.09 |
| Reflections used | 9017 | 9956 | 12022 | 9695 | 12594 | 7594 |
| Free R flagged reflections | 361 | 419 | 549 | 402 | 583 | 299 |
| $R_{cryst}$[e] | 0.207 | 0.184 | 0.200 | 0.193 | 0.197 | 0.189 |
| $R_{free}$[f] | 0.281 | 0.229 | 0.250 | 0.251 | 0.234 | 0.256 |
| rmsd bonds (Å) | 0.007 | 0.008 | 0.009 | 0.006 | 0.010 | 0.007 |
| rmsd angles (˚) | 1.541 | 1.550 | 1.619 | 1.424 | 1.626 | 1.482 |
| Ramachandran plot | | | | | | |
| Most favored (%) | 99.12 | 98.23 | 100 | 100 | 100 | 98.23 |
| Additionally allowed (%) | 0.88 | 1.77 | 0.00 | 0.00 | 0.00 | 1.77 |
| Mean B-factor (Å$^2$) | 62.225 | 50.720 | 49.773 | 52.913 | 48.775 | 59.148 |

[a]Values for the last resolution shell are in parentheses.

[b]$R_{merge} = \Sigma_{hkl}\Sigma_i|I_i(hkl)-\langle I(hkl)\rangle| / \Sigma_{hkl}\Sigma_i I_i(hkl)$, where I(hkl) is the intensity of reflection hkl

[c]$R_{meas} = \Sigma_{hkl}(n/(n-1))^{1/2}\Sigma_i|I_i(hkl)-\langle I(hkl)\rangle| / \Sigma_{hkl}\Sigma_i I_i(hkl)$

[d]$R_{pim} = \Sigma_{hkl}(1/(n-1))^{1/2}\Sigma_i|I_i(hkl)-\langle I(hkl)\rangle| / \Sigma_{hkl}\Sigma_i I_i(hkl)$

[e]$R_{cryst} = \Sigma_{hkl} ||F_{obs}|-|F_{calc}|| / \Sigma|_{Fobs}|$

[f]$R_{free}$ is the cross-validation R-factor computed for the test set of unique reflections.

| | 7GAO | 7GAP | 7GAQ | 7GAR | 7GAS |
|---|---|---|---|---|---|
| **Data collection**[a] | | | | | |
| Wavelength (Å) | 0.92124 | 0.92124 | 0.92124 | 0.92124 | 0.92124 |
| Space group | P43 | P43 | P43 | P43 | P43 |
| Resolution (Å) | 30.56 – 1.69 | 30.63 – 1.68 | 30.69 – 2.14 | 30.58 – 2.07 | 30.66 – 1.91 |
| Last resolution shell (Å) | 1.72 – 1.69 | 1.71 – 1.68 | 2.20 – 2.14 | 2.13 – 2.07 | 1.96 – 1.91 |
| Unit cell parameters | | | | | |
| a,b,c (Å) | 61.07, 61.07, 35.30 | 61.21, 61.21, 35.38 | 61.18, 61.18, 35.47 | 61.03, 61.03, 35.29 | 60.85, 60.85, 35.50 |
| α, β, γ (°) | 90, 90, 90 | 90, 90, 90 | 90, 90, 90 | 90, 90, 90 | 90, 90, 90 |
| Total number of observations | 201357 (10272) | 207746 (10646) | 98511 (7335) | 109090 (8464) | 140622 (9796) |
| Unique reflections | 14738 (749) | 15116 (760) | 7415 (532) | 8083 (600) | 10191 (681) |
| Mosaicity (˚) | 0 | 0 | 0 | 0 | 0 |
| Multiplicity | 13.7 (13.7) | 13.7 (14.0) | 13.3 (13.8) | 13.5 (14.1) | 13.8 (14.4) |
| Mean I/σ(I) | 10.2 (0.5) | 9.9 (0.3) | 6.5 (0.7) | 8.6 (0.3) | 10.8 (0.3) |
| Completeness (%) | 99.9 (99.1) | 99.6 (98.8) | 100.0 (100.0) | 99.8 (98.8) | 99.2 (98.2) |
| $R_{merge}$[b] | 0.262 (13.352) | 0.153 (5.552) | 0.377 (4.262) | 0.173 (7.250) | 0.141 (7.310) |
| $R_{meas}$[c] | 0.273 (13.865) | 0.159 (5.761) | 0.392 (4.426) | 0.181 (7.521) | 0.146 (7.580) |
| $R_{pim}$[d] | 0.074 (3.716) | 0.043 (1.530) | 0.107 (1.187) | 0.050 (1.991) | 0.040 (1.989) |
| **Refinement** | | | | | |
| Resolution (Å) | 30.56 – 1.69 | 30.63 – 1.68 | 30.69 – 2.14 | 30.57 – 2.07 | 30.68 – 1.91 |
| Reflections used | 14048 | 14358 | 7114 | 7677 | 9624 |
| Free R flagged reflections | 660 | 674 | 289 | 304 | 400 |
| $R_{cryst}$[e] | 0.192 | 0.203 | 0.188 | 0.191 | 0.219 |
| $R_{free}$[f] | 0.218 | 0.244 | 0.244 | 0.264 | 0.276 |
| rmsd bonds (Å) | 0.007 | 0.008 | 0.008 | 0.007 | 0.008 |
| rmsd angles (˚) | 1.459 | 1.494 | 1.509 | 1.507 | 1.549 |
| Ramachandran plot | | | | | |
| Most favored (%) | 100 | 99.12 | 100 | 97.35 | 99.12 |
| Additionally allowed (%) | 0.00 | 0.88 | 0.00 | 1.77 | 0.88 |
| Mean B-factor (Å$^2$) | 35.080 | 35.028 | 43.990 | 62.764 | 54.217 |

[a]Values for the last resolution shell are in parentheses.

[b]$R_{merge} = \Sigma_{hkl}\Sigma_i|I_i(hkl)-\langle I(hkl)\rangle| / \Sigma_{hkl}\Sigma_i I_i(hkl)$, where I(hkl) is the intensity of reflection hkl

[c]$R_{meas} = \Sigma_{hkl}(n/(n-1))^{1/2}\Sigma_i|I_i(hkl)-\langle I(hkl)\rangle| / \Sigma_{hkl}\Sigma_i I_i(hkl)$

[d]$R_{pim} = \Sigma_{hkl}(1/(n-1))^{1/2}\Sigma_i|I_i(hkl)-\langle I(hkl)\rangle| / \Sigma_{hkl}\Sigma_i I_i(hkl)$

[e]$R_{cryst} = \Sigma_{hkl} ||F_{obs}|-|F_{calc}|| / \Sigma|_{Fobs}|$

[f]$R_{free}$ is the cross-validation R-factor computed for the test set of unique reflections.
